# Genome-Scale CRISPR Screening Identifies Novel Human Pluripotent Gene Networks

**DOI:** 10.1101/323436

**Authors:** Robert J. Ihry, Max R. Salick, Daniel J. Ho, Marie Sondey, Sravya Kommineni, Steven Paula, Joe Raymond, Elizabeth Frias, Kathleen A. Worringer, Carsten Russ, John Reece-Hoyes, Bob Altshuler, Ranjit Randhawa, Zinger Yang, Gregory McAllister, Gregory R. Hoffman, Ricardo Dolmetsch, Ajamete Kaykas

## Abstract

Human pluripotent stem cells (hPSCs) generate a wide variety of disease-relevant cells that can be used to improve the translation of preclinical research. Despite the potential of hPSCs, their use for genetic screening has been limited because of technical challenges. We developed a renewable Cas9/sgRNA-hPSC library where loss-of-function mutations can be induced at will. Our inducible-mutant hPSC library can be used for an unlimited number of genome-wide screens. We screened for novel genes involved in 3 of the fundamental properties of hPSCs: Their ability to self-renew/survive, their capacity to differentiate into somatic cells, and their inability to survive as single-cell clones. We identified a plethora of novel genes with unidentified roles in hPSCs. These results are available as a resource for the community to increase the understanding of both human development and genetics. In the future, our stem cell library approach will be a powerful tool to identify disease-modifying genes.

**VISUAL ABSTRACT:** 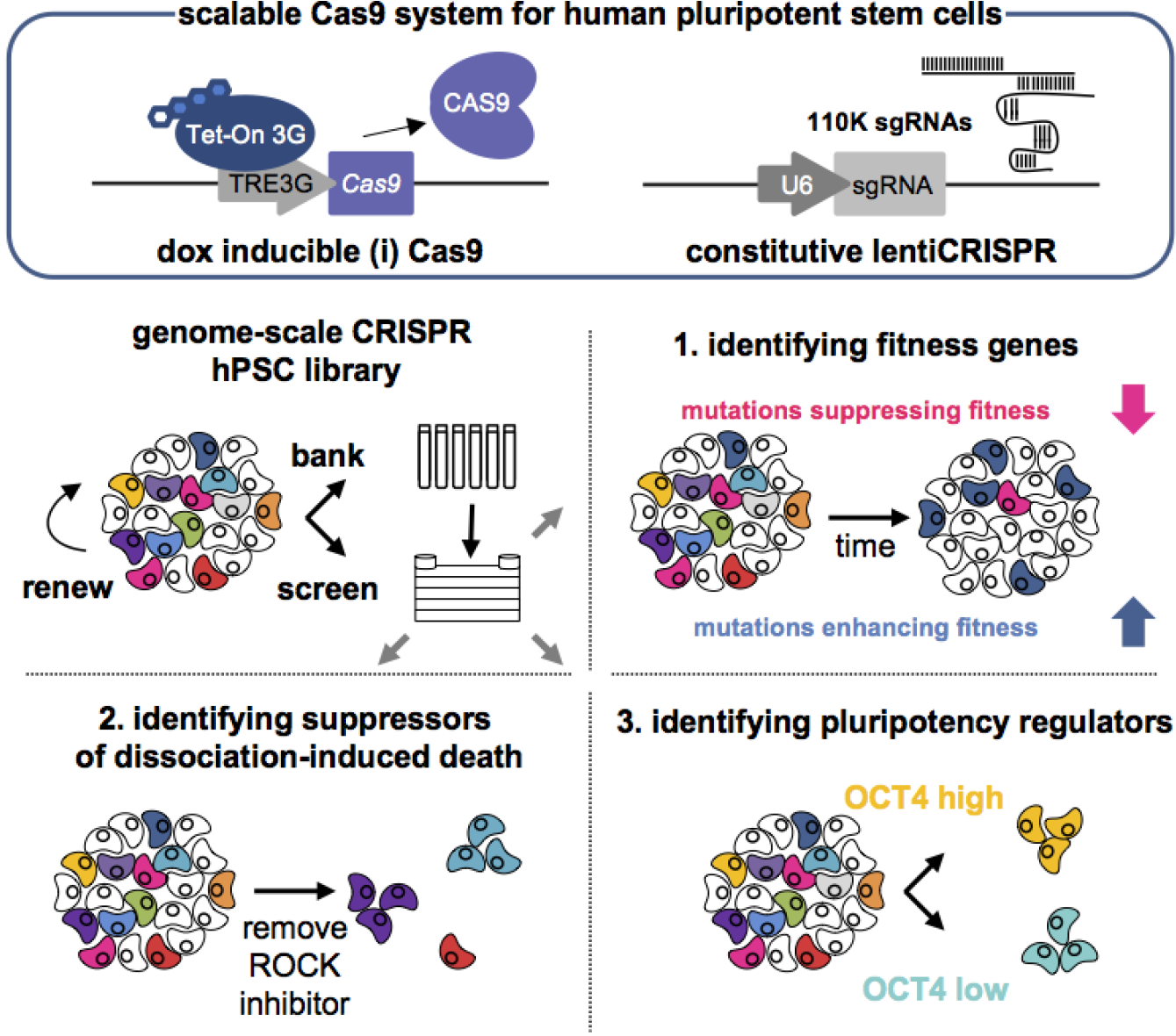

## IN BRIEF

Ihry et al. develop a stable CRISPR/Cas9 genetic screening platform tailored for use in hPSCs that enables unbiased genome-scale genetic screening. The platform exhibits high performance at genome-scale and accurately detects the dropout of core essential genes. Furthermore, proof-of-concept screens exploited hPSC-specific phenotypes to identify regulators of fitness, survival after single-cell dissociation, and pluripotency.

## HIGHLIGHTS

➣ A universal and scalable genetic platform in hPSCs for general use across all lineages
➣ Robust knockout efficiencies translate into high performance screening at genome-scale
➣ Identified stem cell-specific components of *TP53* and *OCT4* genetic networks in hPSCs
➣ Phenotypic screen identifies novel actin/myosin genes regulating membrane blebbing
➣ Identified novel gene sets regulating pluripotency and discovered the roles of *PMAIP1* and *PAWR* in sensitivity to DNA damage and single-cell dissociation

## INTRODUCTION

Human pluripotent stem cells (hPSCs) can be used to generate a wide variety of disease relevant cell types and have the potential to improve the translation of preclinical research by enhancing disease models. Despite the huge potential, genetic screening using hPSCs has been limited by their expensive and tedious cell culture requirements (Chen et al., 2011), and reduced genetic manipulation efficiencies (Ihry et al., 2018). Only a few shRNA screens have been conducted in hPSCs (Chia et al., 2010; Zhang et al., 2013), however shRNAs have a high level of off targets and do not cause a complete loss of function, which is difficult to interpret (DasGupta et al., 2005; Echeverri et al., 2006; Kampmann et al., 2015; McDonald et al., 2017). Currently, the CRISPR/Cas9 system is the genetic screening tool of choice because it can efficiently cause loss-of-function alleles (Cong et al., 2013; Jinek et al., 2012; Mali et al., 2013). Hundreds of genome-scale pooled CRISPR screens have been performed in immortalized human cell lines (Hart et al., 2015; Meyers et al., 2017; Wang et al., 2015). However, in hPSCs the CRISPR/Cas9 system has been primarily used for small-scale genome engineering (Merkle et al., 2015). In genetically intact hPSCs the only genome-scale CRISPR screen to date used methods developed for cancer cells, suffered from technical issues, had poor performance, and identified few developmentally relevant genes (Hart et al., 2014; Shalem et al., 2014). We addressed these technical issues by systematically tailoring the CRISPR/Cas9 system for hPSCs (Ihry et al., 2018). We developed a doxycycline (dox) inducible Cas9 (iCas9) hPSC line and stably infected it with a genome-scale sgRNA library. We banked and expanded the CRISPR-infected hPSC library in the absence of editing (-dox), which enabled us to generate a renewable stem cell pool with stable but inactive sgRNAs. This allowed us to conduct multiple independent screens with the same cell library.

In the first screen, we identified genes that suppress or enhance hPSC fitness over long-term culture. While previous screens have generated gold standard gene lists of core essential genes that reduce cell survival when mutated, little is known about the mutations that enhance survival and proliferation. Unlike core essential genes, these enhancing mutations appear to be cell type-specific and no consistent lists exist for this type of gene (Hart et al., 2014). In hPSCs karyotypic analysis has detected recurrent copy number variations (CNVs) that confer a growth advantage (Amps et al., 2011; Laurent et al., 2011); however, these studies lack gene level resolution. Recently, next-generation sequencing of hundreds of hPSCs identified the recurrence of dominant-negative *TP53* mutations that can expand within a population of hPSCs (Merkle et al., 2017). We mined our data for gene knockouts that enriched in culture and identified many genes, including components of the *TP53* pathway and other known tumor suppressors. We validated the strongest hit, *PMAIP1/NOXA,* which appears to be a stem cell-specific gene conferring sensitivity to DNA damage downstream of *TP53.*

In the second screen, we identified genes required for single-cell cloning. hPSCs have a poor survival rate after dissociation to single-cells, which is detrimental for genome engineering. Multiple groups have extensively characterized death induced by single-cell cloning and have demonstrated the process is similar to but distinct from anoikis and is triggered by a ROCK/Myosin/Actin pathway (Chen et al., 2010; Ohgushi et al., 2010). To prevent death, hPSCs are passaged as clumps or treated with ROCK inhibitors (Watanabe et al., 2007). By subjecting our hPSC mutant library to single-cell dissociation without ROCK inhibitors, we selected for mutations that survive single-cell cloning. sgRNAs for the ROCK and myosin pathways were enriched in the surviving clones. The most enriched gene was the pro-apoptotic regulator *PAWR* (Burikhanov et al., 2009). Validation studies confirmed a novel role for *PAWR* as a component of the actin-cytoskeleton that induces membrane blebbing and cell death caused by single-cell cloning. The additional novel genes identified here will further our understanding about the sensitivity of hPSCs to single-cell cloning.

In the final screen, we utilized a FACS-based OCT4 assay to identify regulators of pluripotency and differentiation. Pluripotency is a defining feature of hPSCs and it allows them to differentiate into all three germ layers. *OCT4/POU5F1*, *NANOG* and *SOX2* are critical transcription factors that maintain pluripotency *in vivo* and *in vitro* (Chambers et al., 2007; Masui et al., 2007; Nichols et al., 1998). *OCT4* and *SOX2* overexpression is commonly used to reprogram somatic cells towards the pluripotent state (Takahashi and Yamanaka, 2006; Takahashi et al., 2007; Yu et al., 2007). By isolating mutant cells with high or low OCT4 protein expression we identified many genes involved in maintaining the pluripotent state along with genes involved with induction of differentiation. Overall these lists of genes provide unique insights into the genetic regulation of human development that could only be identified in normal diploid cells that are not transformed or cancerous.

By using a doxycycline inducible Cas9 (iCas9) hPSC line stably infected with a genome-scale lentiCRISPR library, we were able to bank a CRISPR-hPSC library that was renewable and enabled a high number of independent screens with the same starting library. This allows direct comparison between screens and reduces screen to screen variability. We rigorously tested the system and identified genes important for fitness, pluripotency, and single-cell cloning of hPSCs. Herein we provide a resource with detailed methods and all available data including many novel genes that are involved in hPSC biology. This resource will serve as a parts list of genes that are functionally important for the human stem cell state. Furthermore, the gene sets and methods will increase our systematic knowledge of human pluripotent stem cell biology and will enable additional large-scale CRISPR screens in stem cells and their somatic derivatives.

## RESULTS

### iCas9 system is a self-renewing resource enabling successive genome-wide genetic screens in hPSCs

We set out to develop a high-throughput CRISPR/Cas9 platform for hPSCs which would enable successive rounds of screening from a stable library of lentiCRISPR-infected hPSCs (Fig. 1A, S1). Generating a genome-scale lentiCRISPR hPSC library would enable both the rigorous testing of CRISPR screen performance and the identification of cell type-specific regulators of the pluripotent state. In our previous work, we developed an all-in-one doxycycline (dox) inducible Cas9 (iCas9) transgene that was inactive in the absence of dox (Ihry et al., 2018). The tight control over Cas9 expression allowed us to transduce cells with lentiviruses expressing sgRNAs (lentiCRISPRs) in the absence of dox without causing on-target indels. We tested if it was possible to bank a genome-scale lentiCRISPR infected cell library (5 sgRNAs per gene, 110,000 total sgRNAs) prior to Cas9 mutagenesis (- dox). After one freeze-thaw cycle, NGS analysis revealed no bottlenecking of the library demonstrating the feasibility of banking a large lentiCRISPR hPSC library for repeated screens (Fig. 1B).

### Evaluating the performance of CRISPR screening in iCas9 hPSCs

Next, we performed a fitness screen to evaluate the global performance of the system (Fig. 1C). We benchmarked the performance of the screen by utilizing annotated lists of core essential genes. Core essential genes are required for the survival of all cells; the corresponding CRISPR knockout causes the sgRNAs to be depleted (Hart et al., 2014, 2015). Genome-scale CRISPR screening in hPSCs has been challenging (Hart et al., 2014; Shalem et al., 2014). hPSCs have a strong DNA damage response (DDR) and Cas9-induced double strand breaks (DSBs) cause a significant cell loss (Ihry et al., 2018). Failure to account for Cas9-induced cell loss is problematic for pooled screening because it is critical to maintain representation of each sgRNA-barcoded cell. Our previous work demonstrated a range of Cas9-induced cells loss between 3 to 10-fold across many sgRNAs (Ihry et al., 2018). To prevent bottlenecking of the sgRNA library we conducted the screen in hPSCs at an average of 1000 cells per sgRNA (a total of 110 million infected cells). By doing this we maintained about 4-fold more cells than a typical cancer screen (Hart et al., 2015). During the fitness screen, DNA was sampled before and after dox exposure at days 0, 8, 14, and 18 (Fig. S1). To provide a qualitative measurement of screen performance, we plotted the p-values calculated by the redundant siRNA activity (RSA) test against Q1 based z-scores for a set of core essential and non-essential genes (Hart et al., 2014; König et al., 2007). Before dox treatment the nonessential and core essential genes are randomly distributed within a tight cluster (Fig. 1D). After 18 days of Cas9 treatment the distribution spreads and the essential genes significantly dropout while the non-essential genes remain constant (Fig. 1D, File S1-2).

To quantify performance, we employed the Bayesian Analysis of Gene EssentiaLity (BAGEL) algorithm which calculates a Bayes factor for each gene by determining the probability that the observed fold change for a given gene matches that of known essential genes (Hart and Moffat, 2016).This generates a ranked list of Bayes factors for each gene, which can then be used to quantify screen performance by precision versus recall analysis. In a high-performance screen, essential genes have high Bayes factor scores and the precision versus recall curve gradually drops off as analysis of the ranked list is completed. In contrast, a poor performing screen has a precision versus recall curve that rapidly drops off, indicating many false positives (non-essential genes) with high Bayes factor scores. The sample without dox exposure (untreated) has a randomly ranked Bayes factor list with non-essential and essential genes interspersed and exhibits a poor precision versus recall curve (Fig. 1E). In the day 18 Cas9 (+dox) treated samples, essential genes and non-essential genes segregate from each other and generates a high-performing precision versus recall curve that gradually drops off (Fig. 1E, File S3).

After 18 days of Cas9 exposure, we identified 770 fitness genes at a 5% false discovery rate based on of the precision calculation (Fig. 1F, File S4). Comparing the set of 770 hPSC fitness genes to 1580 core essential genes from cancer lines revealed an overlap of 405 genes (Fig. 1G) (Hart et al., 2015). The remaining 365 specifically dropped out in hPSCs. Both the core and hPSC-specific essential genes are abundantly expressed in hPSCs, further supporting that they are required to maintain hPSCs in culture (Fig. 1H). Our fitness screen in hPSCs correctly identifies the dropout of core-essential genes with accuracy that is on par with CRISPR screens conducted in cancer cell lines. This demonstrates that cancer cells and stem cells share a common set of core essential genes that can be used to benchmark performance. By properly accounting for cell loss caused by Cas9 activity, we have overcome a significant technical barrier that has thwarted previous attempts at genome-scale screening in hPSCs (Hart et al., 2014; Shalem et al., 2014). This demonstrates it is possible to conduct an effective genome-scale CRISPR screen in hPSCs using the methods described here.

### *TP53* pathway mutations specifically enrich during CRISPR fitness screen in hPSCs

Curated lists of genes that enhance fitness during a CRISPR screen do not exist, making it difficult to benchmark the enrichment results (Hart et. al., 2015). By comparing the top ~1000 depleted (RSA-down< −2.25, Files S2) and enriched (RSA-up < −2.25, File S5) genes we observed that 31.8% (301 of 946) of the enriched genes were located on the X and Y chromosomes (H1-hESCs XY, Fig. S3). In contrast, the depleted genes were evenly distributed across all chromosomes. It became apparent that allosome targeting sgRNAs were behaving similarly to non-targeting controls which enrich during a CRISPR screen in hPSCs (Ihry et al., 2018). We observed that sgRNAs causing a single DSB on the X chromosome are less toxic relative to sgRNAs inducing 2 DSBs at the *MAPT* locus despite being able to efficiently induce indels (Fig. S2). sgRNAs targeting genomic amplifications in cancer cell lines exhibit a strong depletion irrespective of the gene targets (Aguirre et al., 2016; Meyers et al., 2017; Munoz et al., 2016). Unlike cancer cell lines, H1-hESCs with a normal diploid karyotype and are very sensitive to DNA damage making the difference between 1 and 2 DSBs significant. After recognizing that the enrichment of sgRNAs on the X/Y chromosomes was related to DSB sensitivity and copy number differences in male H1-hESCs we focused on autosomal genes. In the remaining list of 645 autosomal genes that were enriched (RSA-up −2.25) we identified 41 tumor suppressor genes (Zhao et al., 2016).

The second most enriched gene was *TP53,* which confirms the selective pressure imposed by Cas9-induced DSBs in hPSCs during a CRISPR screen (Fig. 2A). Consistent with this *TP53* mutants are able to suppress cell loss induced by Cas9 activity (Ihry et al., 2018). Throughout the 18-day screen, the representation of sgRNAs targeting *TP53* (Chr. 17), the checkpoint kinase, *CHEK2* (Chr. 22), and the proapoptotic regulator, *PMAIP1* (Chr. 18), increased in a time-dependent manner (Fig. 2B). Database mining for associations with *TP53* among the enriched genes identified 20 genes with direct connections to *TP53* (Fig. 2C, File S6). 19 of which are expressed in H1-hESC and are on autosomal chromosomes. Although *EDA2R* is on the X chromosome, it was included because its mRNA increases in response to DNA damage in hPSCs (Ihry et al., 2018). We hypothesized that these genes could include additional regulators responsible for the extreme sensitivity to DNA damage in hPSCs.

*PMAIP1* was the most enriched gene in the screen. PMAIP1 has been implicated in TP53-dependent cell death and functions by sensitizing cells to apoptosis by antagonizing the anti-apoptotic protein MCL1 at the mitochondria (Kim et al., 2006; Perciavalle et al., 2012; Ploner et al., 2008). *PMAIP1* is highly expressed in hPSCs and its expression marks the pluripotent state (Mallon et al., 2013). We confirmed this by examining *PMAIP1* expression in 2 iPSC and 2 hESC lines. Analysis of RNA-seq experiments confirmed that *PMAIP1* was highly expressed during the pluripotent state and drops during neuronal differentiation using the *NGN2* transgene (Fig. 3A). Additionally, GTEx data revealed that *PMAIP1* has a highly restricted expression pattern and is not expressed in most tissues (GTEx Analysis Release V7). Although it has been demonstrated that *PMAIP1* expression is maintained by *OCT4* in testicular germ cell tumors (Gutekunst et al., 2013), no functional connection has been made in hPSCs. We tested if *PMAIP1* expression was maintained by the pluripotency network by differentiating cells or knocking out *OCT4.* Under these conditions qPCR detected a significant decrease in *PMAIP1* mRNA (Fig. 3B). In thymocytes, *PMAIP1* mRNA is induced by TP53 (Khandanpour et al., 2013). We knocked out *TP53* in hPSCs but did not detect a reduction in *PMAIP1* mRNA (Fig 3B). hPSCs constitutively express high levels of *PMAIP1,* and DNA damage does not increase *PMAIP1* expression, further supporting *PMAIP1* mRNA as pluripotency-dependent and TP53-independent (Ihry et al., 2018).

Prior work demonstrated that cancer cell lines have a reduced DNA damage response relative to hPSCs (Ihry et al., 2018). Consistent with this we did not observe an enrichment of *PMAIP1* sgRNAs in 14 independent CRISPR screens conducted in cancer cell lines despite using libraries containing the same PMAIP1-targeting sgRNAs (Fig. 3C). This suggested that *PMAIP1* is responsible for making hPSCs sensitive to DNA damage. To test the functional consequences of *PMAIP1* mutations, we knocked out *PMAIP1* in the iCas9 cell line using transient exposure to synthetic crRNAs (Fig. S3). We tested if *PMAIP1* mutants were resistant to DSB-induced death by using lentiCRISPRs to deliver a sgRNA targeting *MAPT*, a neuronal gene not expressed in hPSCs. In the absence of Cas9 (-dox), both control and *PMAIP1* mutant iCas9 cells grow at a similar rate while expressing an sgRNA. In the presence of Cas9 (+dox) and a sgRNA, control cells die while *PMAIP1* mutants are able to survive despite efficient DSB induction (Fig. 3D, S3, S4E). qPCR analysis of the TP53-target genes *P21* and *FAS* detected an elevated expression in *PMAIP1* mutants compared to controls (Fig. 3E). Despite having an active *TP53, PMAIP1* mutants survive. This indicates *PMAIP1* is downstream of *TP53* activation and is consistent with its known role as a sensitizer to apoptosis (Ploner et al., 2008). In hPSCs cell death is the predominant response to DNA damage. Unlike *PMAIP1* mutants, *P21* mutants (>80% indels) are unable to suppress DSB-induced toxicity (Fig. S4E), and *P21* sgRNAs did not enrich in H1-hESCs. In contrast, in a genome-scale screen in retinal pigment epithelial cells, where Cas9 activity causes TP53-dependent cell cycle arrest rather than apoptosis, sgRNAs targeting *TP53* and *P21/CDKN1A* were enriched while *PMAIP1* sgRNAs were not (File S5) (Haapaniemi et al., 2017). Overall, these results indicate that *PMAIP1* plays a role in the sensitivity of hPSCs to DNA damage and highlights the ability of genome-scale CRISPR screens to identify cell type-specific genes important for the fitness of pluripotent stem cells.

### Genetic screen for suppressors of dissociation-induced death

We next tested our ability to identify phenotypic regulators of human developmental processes. Human PSCs, unlike mouse PSCs, are very sensitive to dissociation and die in the absence of ROCK-inhibitors (Ohgushi et al., 2010). During dissociation of hPSCs, Rho and ROCK become activated. This leads to the phosphorylation of myosin, which causes membrane blebbing and cell death. To promote survival, inhibitors that target ROCK (Y-27632/thiazovivn) or myosin (blebbistatin) are used routinely during hPSC passaging (Chen et al., 2010; Watanabe et al., 2007). Very few cells survive dissociation in the absence of ROCK inhibitors. Importantly, this phenotype is developmentally rooted. hPSCs are epiblast-like and cells that fail to incorporate into the polarized epithelium of the epiblast undergo cell death in the embryo (Ohgushi et al., 2010). To gain a deeper understanding of the genes involved, we screened for suppressors of dissociation-induced death.

During the day 14 passage of the fitness screen we plated an additional replicate of the genome-scale mutant cell library in the absence of thiazovivin (Fig. 4A, S1). The majority of the cells died during this process and the surviving cells were maintained for two weeks until large colonies were visible, at which point DNA was isolated and sgRNAs sequences were recovered by NGS. We identified 76 genes with 2 or more independent sgRNAs present in cells that survived dissociation without thiazovivin (Fig. 4C and File S7). As expected, multiple sgRNAs targeting *ROCK1* and *MYH9,* the genetic targets of ROCK inhibitors and blebbistatin, were recovered. Myosin is a hexameric motor protein that is comprised of 6 subunits with 3 subtypes. The screen recovered 3 sgRNAs for *MYH9,* a myosin heavy chain, 5 sgRNAs for *MYL6,* a non-phosphorylated myosin light chain, and 2 sgRNAs for the ROCK target *MYL9/MLC2,* a phosphorylated myosin light chain. There are many myosin proteins, but our screen identified 3 out of 5 of the most abundantly expressed in hPSCs (Fig. 4B). This reiterates the importance of myosin activation in membrane blebbing and dissociation-induced death. A number of additional genes have roles in the actin/myosin network or cytoskeleton, including *DAPK3, PAWR, OPHN1, FLII* and *KIF3A* (Fig. 4D). Overall STRING-DB analysis detected a connected set of genes with ties to the actin/myosin regulatory network (Fig. 4D). In general members of this network did not enrich during the fitness screen suggesting they specifically regulate dissociation-induced death and not fitness (Fig. 4E).

### *PAWR* is required for dissociation-induced death

For follow-up studies, we focused on the strongest hit from the screen, the pro-apoptotic regulator *PAWR* (Hebbar et al., 2012). *PAWR* has no known biological role in the early embryo or during dissociation-induced death of hPSCs. The screen recovered all 5 sgRNAs targeting *PAWR* in the genome-scale library and the barcode reads were highly abundant (Fig. 4C). Unlike sgRNAs for the *TP53* pathway that enrich throughout the CRISPR screen, *PAWR* and *MYL6* have no effect on fitness in the presence of thiazovivin (Fig. 4E). We repeated the results using 3 independent lentiCRISPRs to knock out *PAWR* in iCas9 expressing H1-hESC cells. *PAWR* mutants are able to survive without thiazovivin while control cells do not (Fig. S4). To independently validate these results, we used CRISPR mediated interference (CRISPRi) to knockdown the expression of *PAWR* mRNA without causing DNA damage or genetic mutations (Qi et al., 2013). By using a H1-hESC line constitutively expressing dCas9 fused to a KRAB domain and sgRNAs targeting *PAWR* promoter we also detected an increase in the survival of dissociated cells in the absence of thiazovivin (Fig. S4).

To conduct detailed analysis of *PAWR* mutants we exposed cells to Cas9 RNPs targeting *PAWR* and generated a knockout cell line with a normal karyotype (Fig. S4). The suppression of dissociation-induced death is specific to *PAWR* mutants, as *PMAIP1* mutants are unable to survive passaging without thiazovivin (Fig. 5A-B). Conversely, *PAWR* mutants are unable to survive DSB-induced toxicity, further demonstrating the specificity of the respective phenotypes (Fig. S4E). We next examined how *PAWR* mutants survive by time-lapse microscopy. After single cell dissociation, control cells without thiazovivin exhibit membrane blebbing and subsequently die (Fig 5C). Conversely, *PAWR* mutants without thiazovivin have greatly reduced blebbing and survive as single cells by extending cell projections, which promote attachment and survival (Fig 5C). We further examined cytoskeletal organization using phalloidin to stain filamentous (F-) ACTIN. The thiazovivin-treated cells have an increased surface area, a fanned-out shape with actin stress fibers, and a large circular adhesion belt-like structure (Fig. 5C). In the absence of thiazovivin, control cells have many small actin rings which mark membrane blebs. *PAWR* mutants without thiazovivin exhibit reduced membrane blebbing and small actin rings (Fig. 5C).

Molecularly, *PAWR* has dual roles as a transcriptional repressor that causes cell death or as an actin binding protein that regulates contractility (Burikhanov et al., 2009; Johnstone et al., 1996; Vetterkind and Morgan, 2009). Despite having abundant *PAWR* mRNA, PAWR protein is post-trancriptionally regulated and induced by dissociation in hPSCs (Fig. S4-5). Immunofluorescence did not detect PAWR protein in the nucleus; however, we did observe localization of PAWR with F-ACTIN in both thiazovivin treated and untreated cells (Fig. 5D). PAWR localized to adhesion belt-like structures in the presence of thiazovivin and to membrane blebs in the untreated cells after dissociation. Additional hits from the screen, PRKCZ and SCRIB, also localized to membrane blebs (Fig. S6). Both PRKCZ and SCRIB are expressed in the early mouse embryo and exhibit a ROCK-dependent cell polarity (Kono et al., 2014). Cumulatively, we have identified a novel role for *PAWR* in dissociation-induced death. *PAWR* mutants survive dissociation without ROCK inhibitors because of a failure to initiate membrane blebbing and downstream caspase activation (Fig S5) (Ohgushi et al., 2010). Furthermore, *PAWR* is a known proapoptotic factor and we have demonstrated that PAWR protein is induced upon dissociation and colocalizes with the ACTIN network that is responsible for initiating membrane blebbing and subsequent cell death.

### FACS-based screen for regulators of pluripotency

Although our fitness screen detected a significant hPSC-specific dropout of *OCT4* other critical regulators of pluripotency like *NANOG* did not affect fitness (Fig 1E, File S4). A genome scale fitness screen in mouse ESCs had similar results and only reported the dropout of three genes regulating blastocyst development (Koike-Yusa et al., 2014). Pluripotency and cellular fitness of hPSCs may not be related and this indicates that some differentiated cell types may not exhibit changes in fitness when cultured in the pluripotent media. To specifically identify regulators of human pluripotency we conducted a FACS-based pooled screen using an OCT4 antibody. One year after conducting the first fitness screen we thawed a genome-scale CRISPR cell library and expanded them prior to conducting the screen (Fig. S1). The cells were mutagenized with Cas9 for 8 days prior to FACS to collect cells with high (OCT4^HIGH^) and low (OCT4^LOW^) OCT4 expression (Fig. 6A). Log2(fold change) was calculated by comparing the OCT4^LOW^ group to the OCT4^HIGH^ group. We plotted the p-values calculated by the RSA test against Q1 and Q3 based z-scores for OCT4^LOW^ and OCT4^HIGH^, respectively (Fig. 7B, File S8). Importantly, we detected a significant enrichment of *OCT4* and *NANOG* targeting sgRNAs in the OCT4^LOW^ group. We also detected an enrichment of *TGFBR1/2* genes required to maintain the culture of hPSCs and the chromatin regulators *EP300* and *SMARCA4/BRG1* which regulate *OCT4* expression and function in hPSCs (Chen et al., 2011; King and Klose, 2017; Singhal et al., 2010; Wang et al., 2012). Using STRING-db we identified a core network of genes connected to *OCT4* in the OCT4^LOW^ group (Fig. 6C). This highlights the ability of phenotypic CRISPR screening to identify relevant gene networks. Additionally, the OCT4^HIGH^ group identified factors that promote differentiation such as *HAND1, KDM5B, and EIF4G2*, (File S8) (Hough et al., 2006; Kidder et al., 2013; Yamanaka, 2000). Many of the genes in these lists have published roles in regulating pluripotency, reprogramming or embryonic development, and further investigation of the less studied genes will reveal novel insights into the human pluripotent state (File S8-9).

### Identification of hPSCs specific fitness and pluripotency gene networks

Extensive CRISPR screening in cancer cell lines has provided a wealth of knowledge about their genetic dependencies. However, the static state of these cells has revealed less about genes with developmental functions (Hart et al., 2015). To compare hPSC results to cancer cell lines we conducted pairwise Pearson correlation coefficients using Bayes factors distributions (cancer data from Hart et al., 2015). The analysis revealed that hPSCs formed a distinct cluster (Fig. 7A) and is consistent with the partial overlap between core essential genes in cancer and fitness genes in hPSCs (52% Fig. 1E). To focus on gene networks specific to hPSCs we conducted bioinformatics analysis comparing 829 core essentials cancer genes (essential for 5/5 cell lines, Hart et al., 2015) to hPSC-specific gene sets identified by our screens. A total of 661 hPSC-specific genes were obtained from the fitness screen (365 depleted and 20 enriched p53 related genes), dissociation-induced death screen (76), and the OCT4 FACS screen (212) (File S10). The gene lists were analyzed using the PANTHER classification system (pantherdb.org). PANTHER pathway analysis identified a greater diversity of 92 enriched pathways in hPSCs and only 38 in the cancer lines (Fig. 7B). In accordance with this we also detected an increase in the number of genes with receptor and signal transducer activity molecular functions (Fig. 7C). The hPSC-enriched pathways included several expected regulators: FGF, TGF-Beta, and WNT (Fig. 7B). FGF2 and TGFβ are critical components of E8 media and are required to maintain pluripotency *in vitro* (Chen et al., 2011). WNT signaling regulates both differentiation and pluripotency in ESCs (Sokol, 2011). Activation of EGF, PDGF, and VEGF is also important for the maintenance of the pluripotent state in (Fig. 7B) (Brill et al., 2009). P53 and CCKR/Rho GTPases pathways, which regulate apoptosis and have critical roles in determining the sensitivity of hPSCs to DNA damage and enzymatic dissociation, were also enriched (Fig. 7B). Examination of the biological processes gene ontology revealed an increase in the number of development and multicellular organism genes (Fig. 7C). Furthermore, subdividing the developmental process category revealed enrichment of genes regulating cell death, differentiation and early developmental stages (Fig. 7C). Globally these results highlight the identification of cell type-specific genes regulating different aspects of the pluripotent state. By screening for regulators of 3 fundamental processes governing the culture of hPSCs we have successfully identified known regulators in addition to a plethora of novel genes involved in stem cell biology (Fig. 7D). These genes are a blueprint for human pluripotency and will serve as a useful resource for the human development and stem cell communities.

## DISSCUSSION

The use of hPSCs in large-scale functional genomics studies has been limited by technical constraints. Prior to our study, it was unclear that genome-scale CRISPR screens were possible in hPSCs as the first attempts had poor performance (Hart et al., 2014; Shalem et al., 2014). We overcame this by building a high-performance 2-component CRISPR/CAS9 system for hPSCs. The performance, across many sgRNAs, made the platform amenable to high-throughput screening. Using iCas9 in hPSCs it is possible to perturb hundreds of genes in arrayed format or the entire genome in pooled format. The system is renewable and a pool of stem cells with sgRNAs to the entire genome can be banked, distributed and utilized for successive screens in hPSCs and their differentiated progeny. Beyond technical proficiency we identified genes that regulate fundamental stem cell processes such as self-renewal, their inherent sensitivity to DNA damage, single-cell cloning, pluripotency and differentiation.

First, we identified 770 genes required for the self-renewal of hPSC. A majority of these genes have established roles in fitness while 365 of these genes are novel and specific to hPSCs. This set of genes could be used to inform a systematic approach to improve the consistency, robustness and user-friendliness of hPSC culture conditions. During the fitness screen, we also determined that Cas9 activity imposes a selective pressure on DNA damage sensitive hPSCs. This caused an enrichment of 20 genes that are connected to *TP53.* Consistent with this, dominant negative *TP53* mutations and deletions recurrently occur and provide a selective advantage during the culture of hPSCs (Amir and Laurent, 2016; Merkle et al., 2017). In addition to *TP53,* we identified a hPSC-specific role for *PMAIP1* in determining the extreme sensitivity of hPSCs to DNA damage. Like *TP53,* deletions of chromosome 18 spanning the *PMAIP1* locus have been recurrently observed during hPSC culture (Amps et al., 2011) and suggest that *PMAIP1* deletion may be responsible for enhanced survival of these lines. These TP53-related genes have the potential to improve the efficiency or safety of genome engineering through transient inhibition or by monitoring their spontaneous mutation rate during hPSC culture (Ihry et al., 2018; Merkle et al., 2017).

Second, we identified 76 genes that enhance the survival of hPSCs during single-cell dissociation. Collectively, the screen uncovered an Actin/Myosin network required for membrane blebbing and cell death caused by dissociation. We identified a novel role for *PAWR,* a pro-apoptotic regulator that is induced upon dissociation. *PAWR* is required for membrane blebbing and subsequent death of dissociated hPSCs in the absence of ROCK inhibitors. Importantly, the results are developmentally relevant, and we also identified *SCRIB* and *PRKCZ* which are known regulators of cell polarity in the early mouse embryo (Kono et al., 2014).

*PAWR* has been shown to physically interact with *PRKCZ* and suggests a potential link between cell polarity and dissociation-induced death (Díaz-Meco et al., 1996). These hits appear to be related to the polarized epiblast-like state of primed hPSC and could explain why polarized hPSCs are sensitive to dissociation whereas unpolarized naïve mESC are not (Takashima et al., 2015). Lastly, this set of genes could enable focused approaches to improve the single-cell cloning efficiencies of hPSCs and the culture of naïve hPSCs.

In the final screen, by subjecting our hPSC CRISPR library to FACS sorting on OCT4 protein, we identified 113 genes that are required to maintain pluripotency and 99 genes that potentially regulate differentiation. Overall, the screen identified an entire network of genes related to OCT4. 19 of these genes have previously indicated roles in pluripotency, embryo development and reprogramming (File S9). Future studies on the novel genes in the list will yield new insights about the genetic control of pluripotency and differentiation. These gene sets could guide rational improvements to protocols for the maintenance, differentiation and reprogramming of hPSCs.

Overall, the results highlight the ability of unbiased genome-scale screens to identify critical and novel regulators of human pluripotent stem cell biology. Future investigation into the genes provided here will be a step toward the genetic dissection of the human pluripotent state. Herein we provide simple and scalable work flows that will lower the entry barrier for additional labs to conduct large-scale CRISPR screens in hPSCs. The scalable and bankable platform described here is a renewable resource that will allow for successive screens and the distribution of CRISPR infected hPSC libraries. The platform could potentially be used to improve the generation, culture and differentiation capacity of hPSCs. It can be applied to the study of development and disease in wide variety of differentiated cell types. Established protocols for neurons, astrocytes, cardiomyocytes, hepatocytes and beta-cells can be exploited to dissect the genetic nature of development and homeostasis in disease relevant cell types. This resource opens the door for the systematic genetic dissection of disease relevant human cells in way that was only before possible in model organisms.

**Figure 1:**
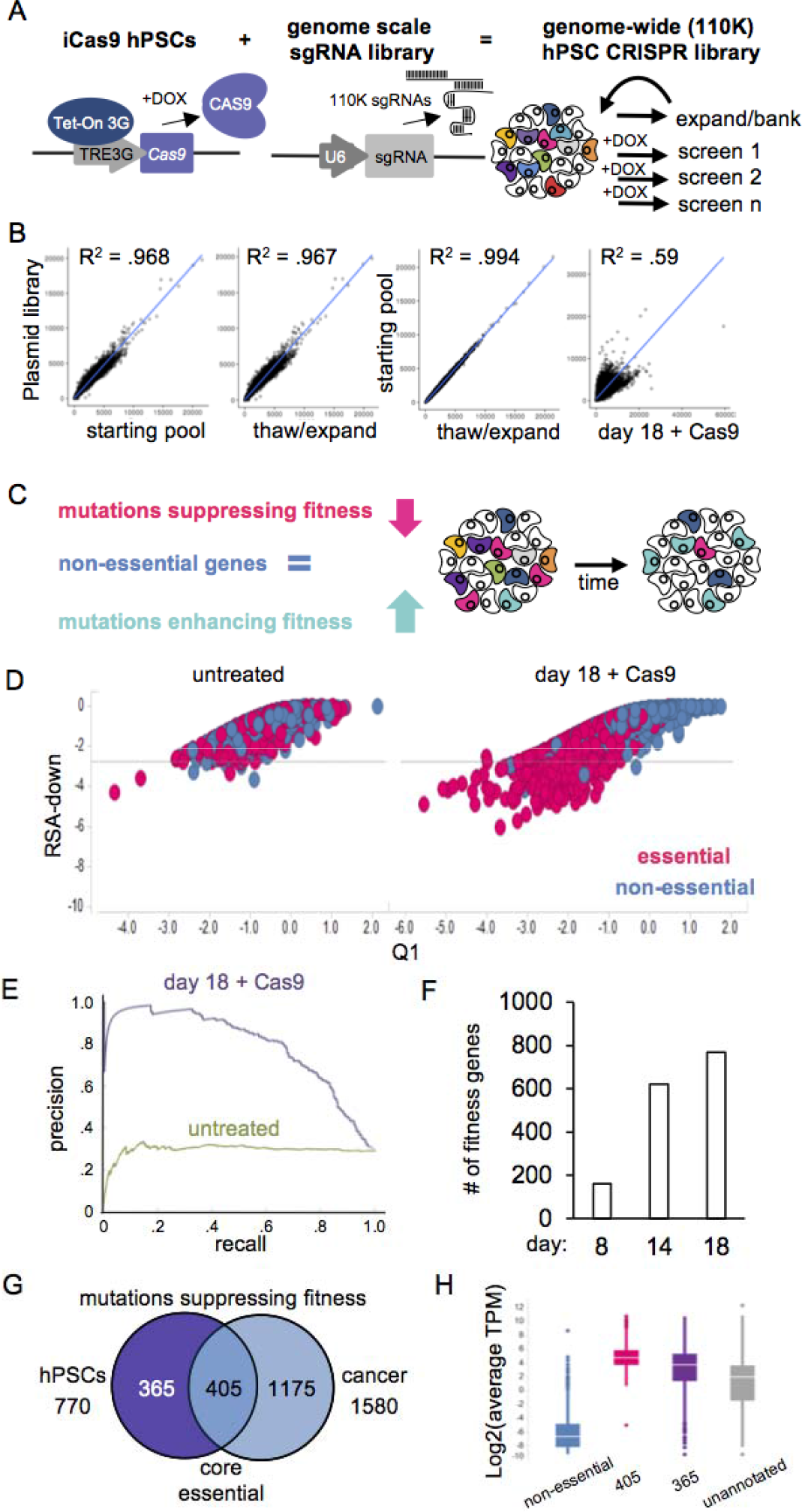
A self-renewing Dox-inducible CAS9/genome-wide sgRNA cell library enables multiple high performance CRIPSR screens in human pluripotent stem cells. (A) Diagram depicting the iCas9 platform for genome-scale CRISPR screening in hPSCs. The iCas9 platform consists of a dox-inducible Cas9 transgene knocked in the AAVS1 locus of H1-hESCs and lentiviral delivery of constitutively expressed sgRNAs. iCas9 hPSCs were transduced (.5 MOI) at scale. After a week of expansion and selection for lentiCRISPRs iCas9 hPSCs were either banked or subjected to Cas9 mutagenesis for screening. (B) Correlation of normalized sgRNA counts reveals that freeze/thaw expanded samples (-dox) have high correlation with the plasmid library and the starting pool of infected iCas9 hPSCs at day 0 of the screen. Day 18 samples have been exposed to dox (+Cas9). (C) Diagram depicting three categories of genes that enrich (enhance), deplete (suppress) or remain constant during a fitness screen. (D) Scatter plot depicting gene level results for core essential (pink) and non-essential genes (blue). Without Cas9 treatment (-dox), cells expressing sgRNAs targeting core essential and non-essential genes are interspersed and have an RSA > −2.75 (marked by dashed line). After 18 days of exposure to Cas9 (+dox), sgRNAs targeting essential genes dropout to less than RSA −2.75. Y-axis is RSA value. X-axis marks the Z-score (Q1). Non-essential and core essential gene list from Hart et. al., 2014. (E) Precision v. recall analysis of genome-scale CRISPR screening data in H1-hESCs. Cas9 expressing cells (purple) exhibit a PR curve that gradually slopes off, whereas cells without Cas9 (green) exhibit a PR curve that immediately decreases. (F) Fitness gene calculation based on 5% FDR on y-axis. Each condition labeled on the x-axis. (G) Venn diagram comparing 770 depleted genes in hPSCs to 1580 core essential genes identified by screening cancer cell lines (Hart et. al., 2015). 405 of the hPSCs depleted genes overlap while the remaining 365 are specifically depleted in pluripotent stem cells. (H) Genes that dropout in CRISPR screen are abundantly expressed. Y-axis depicts the log_2_ transformation of average TPM values from 20 independent RNA-seq experiments in H1-hESC. For each box blot the median is depicted by white line flanked by a rectangle spanning Q1-Q3. X-axis depicts gene categories. In blue are non-essential genes from Hart et al., 2014, in pink are 405 core-essential genes, in purple are 365 stem cell-specific essential genes and in gray are the remaining unannotated genes.

**Figure 2:**
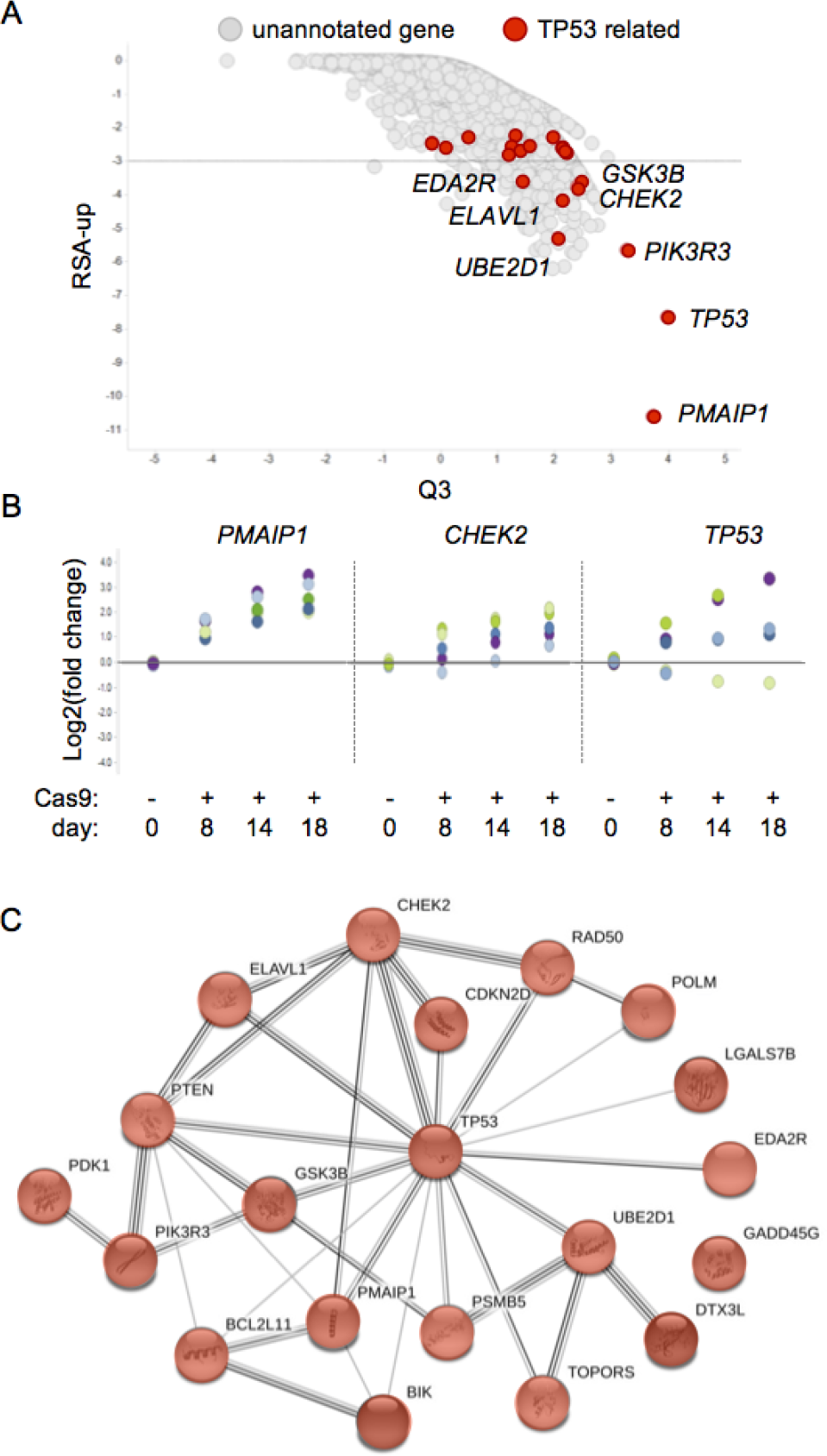
*TP53* pathway mutations enrich during CRISPR screen in hPSCs. (A) Scatter plot depicting gene level results for the genome-scale screen. After 18 days of exposure to Cas9 (+dox) sgRNAs targeting *TP53* related genes enrich during the screen; *PMAIP1 (NOXA)* (RSA - 10.6, Q3 3.7), *TP53* (RSA −7.64, Q3 4), and *CHEK2* (RSA −3.6, Q3 2.4). A total of 302 genes have a RSA score < −3 (marked by dashed line). *TP53* related genes with RSA scores <−2.25 are marked in red. Y-axis is a p-value generated from RSA Up analysis. X-axis marks the Z-score (Q3). (B) Plot showing the time-dependent increase in 5 independent sgRNAs targeting *PMAIP1, CHEK2* and *TP53* during CRISPR screen. NGS quantifies representation of lentiCRISPR-infected cells. Samples were normalized to the day 0 population. Y-axis represents log2 (fold change). Day 0 data shown is from freeze/thaw samples. X-axis plots each condition over time. Cas9 + samples were treated with dox to induce Cas9 expression. (C) Shown are gene knockouts (20) that enrich during CRISPR screen that are connected to *TP53* and play roles in either DNA damage response and apoptosis. 946 enriched genes in RSA up < −2.25 identified by STRING-DB analysis.

**Figure 3:**
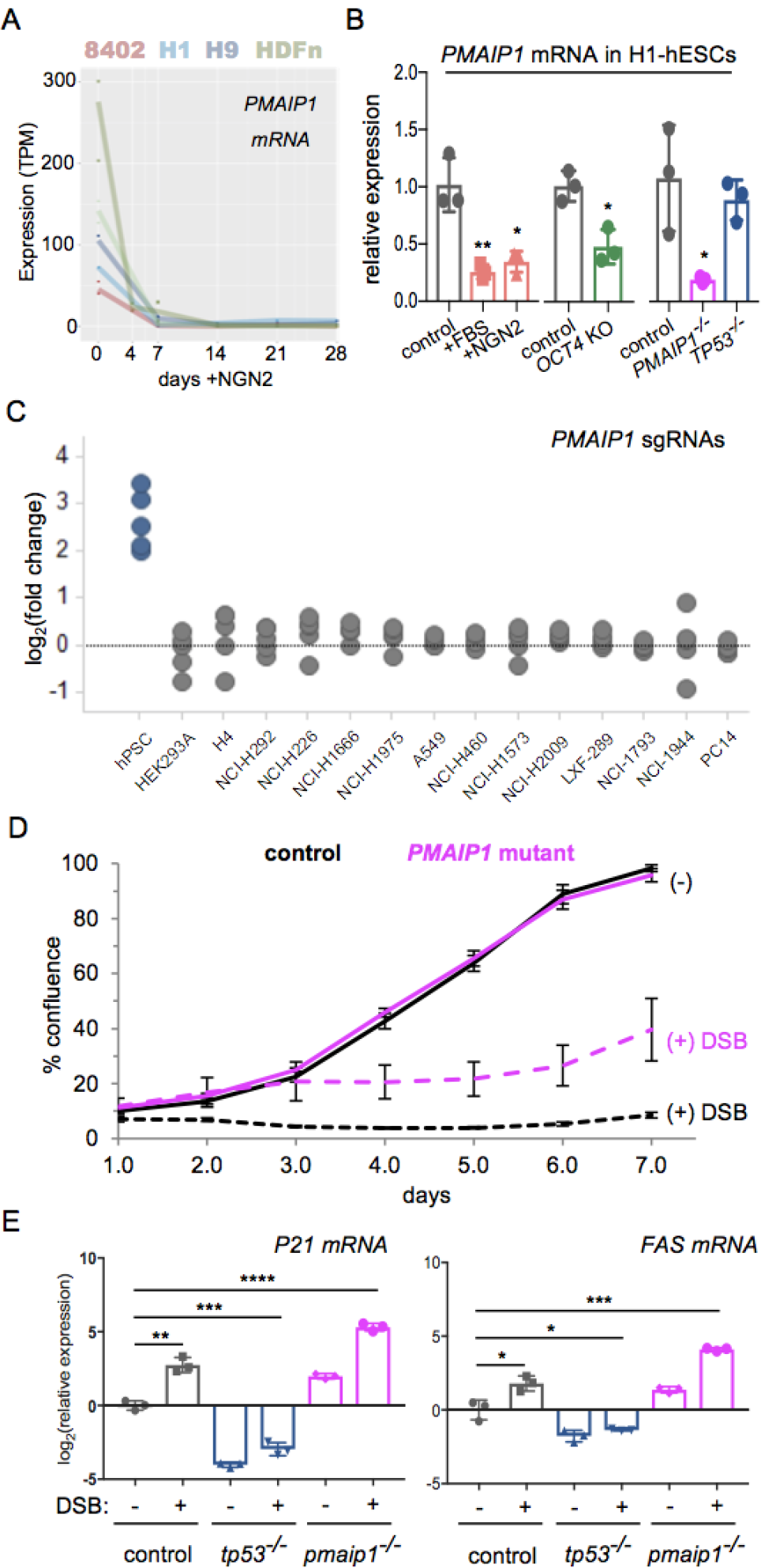
*PMAIP1* confers sensitivity to DNA damage in hPSCs. (A) *PMAIP1* is highly expressed in hESCs and iPSCs. Y-axis represents expression in Transcripts Per Kilobase Million (TPM). H1-hESCs (n=1), H9-hESCs (n=3), 8402-iPSCs (n=3) and HDFn-iPSCs (n=4). X-axis represents days after induction of a doxycycline inducible NGN2 expression cassette. (B) qPCR confirms *PMAIP1* mRNA is dependent on the pluripotent state. Y-axis is relative expression and each bar represents mean relative expression. X-axis is each condition. Control = hPSCs in E8 media, +FBS = 3 days exposure to 10% FBS and DMEM, +NGN2 = 3 days exposure to NGN2, *OCT4* KO = mutant pool after 6 days exposure to iCas9 and sgRNA targeting *OCT4,* PMAIP1−/− = complete knockout cell line, *TP53−/−* = complete knockout cell line. n=3 independent mRNA samples per sgRNA, error bars +/− 1 std. dev. Unpaired two-tailed t-test, equal variances *p<.05, **p<0.01. (C) *PMAIP1* targeting sgRNAs specifically enrich during CRISPR screen in hESC but not cancer cell lines. X-axis plots CRISPR screens conducted in H1-hESC lines and 14 additional transformed lines. 5-independent sgRNAs marked by dots. Y-axis represents log2(fold change). (D) *PMAIP1* mutant hPSCs are insensitive to DNA damage. Live imaging of confluence in *MAPT* sgRNA expressing iCas9 cells +/− DSB (+dox/Cas9) in control or *PMAIP1*^−/−^knockout cell line. Unlike DSB treated control cells the *PMAIP1*^−/−^mutants survive in the presence of DSBs. Black lines indicate control and magenta lines indicate *PMAIP1*^−/−^mutants. Solid lines are without dox and dashed lines are cultured with dox. Y-axis is percent confluency each point represents mean (4 images per well, n=3 wells). error bars +/− 1 std. X-axis time in days of treatment. (E) qPCR of *TP53* target genes indicates *PMAIP1* functions downstream of *TP53. P21 and FAS* mRNA is induced by *MAPT* targeting sgRNAs in iCas9 control cells 2 days after dox treatment. *PMAIP1*mutants exhibit increased levels of *P21* and *FAS* mRNA which is absent in *TP53*mutants. Y-axis is relative expression is calculated by comparing the *MAPT* targeting sgRNA plus (+dox) or minus Cas9 expression (-dox). Unpaired two-tailed t-test, equal variances *p<.05, **p<0.01, ***p<0.001, ****p<0.0001.

**Figure 4:**
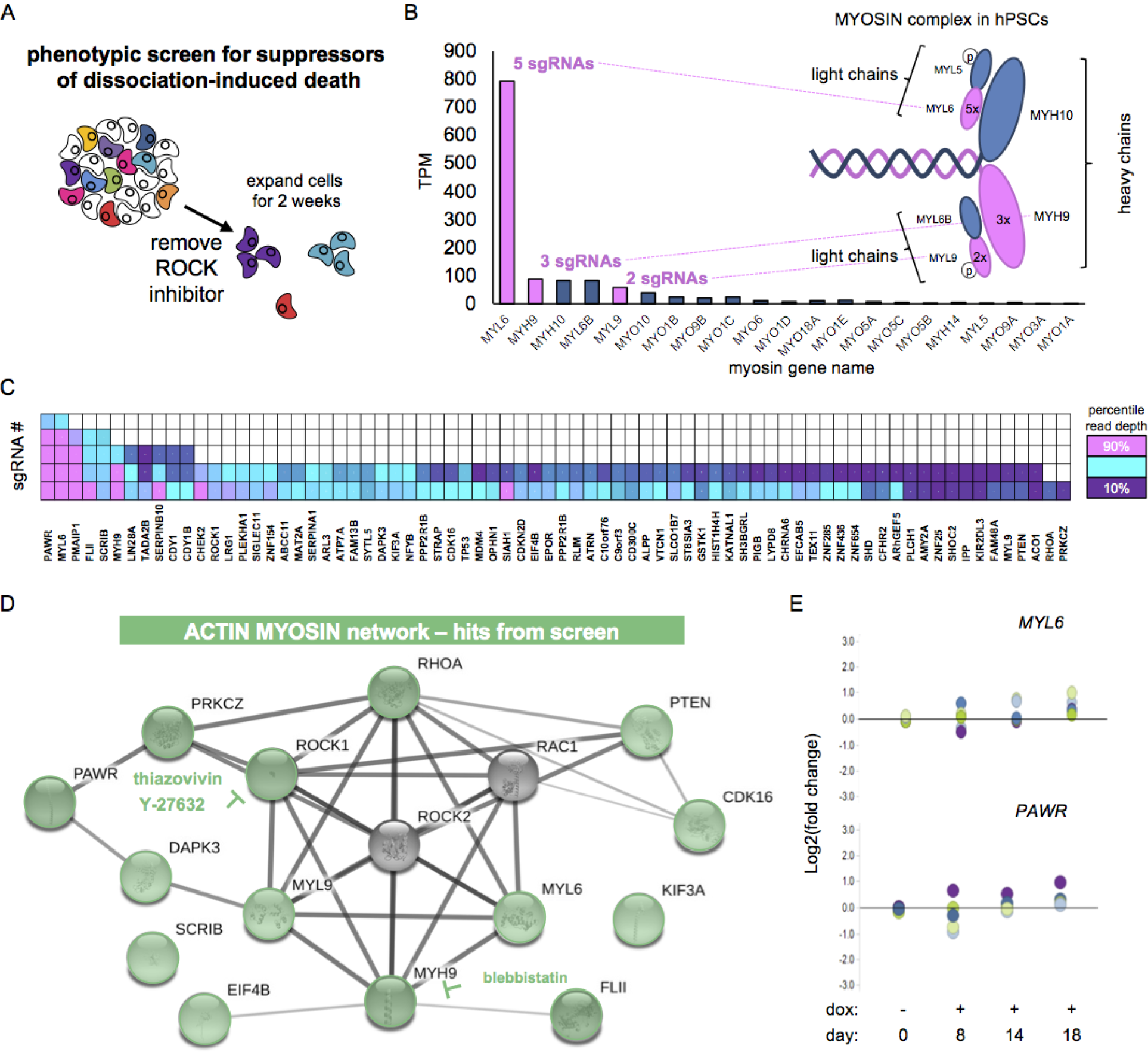
Genetic screen for suppressors of dissociation-induced death. (A) Diagram depicting screen shows the dissociation and replating of the genome-scale mutant cell library without the thiazovivin (ROCK inhibitor). Most cells did not survive the treatment; however, at the end of two weeks some large colonies were recovered for DNA isolation and NGS analysis. (B) The screen recovered 3 out of 6 subunits of the hexameric MYOSIN motor protein that regulates blebbing in hPSCs. Y-axis average TPM in H1-hESC. X-axis myosin genes expressed >1 TPM in H1-hESCs. Myosin genes recovered by screen in pink. Genes that were not detected by screen in blue. (C) Heat map depicting number of sgRNAs recovered on y-axis and each gene on x-axis. Colors indicate the abundance of each sgRNA recovered. Pink marks greater than 90 percentile and purple marks less than 10^th^ percentile. (D) String-DB analysis highlights ACTIN and MYOSIN gene network among mutations that allow cells to survive dissociation in the absence of ROCK inhibitors. Hits from screen marked in green. (E) *MYL6* and *PAWR* specifically regulate survival after dissociation and do not enrich during CRISPR screen. Each dot represents 5 independent sgRNAs per gene and NGS quantifies representation of lentiCRISPRs infected cells. Samples were normalized to the day 0 population and y-axis represents log2(fold change). Day 0 data shown is from freeze/thaw samples maintained at 2100x. X-axis plots each condition over time. Cas9 + samples were treated with dox to induce Cas9 expression.

**Figure 5:**
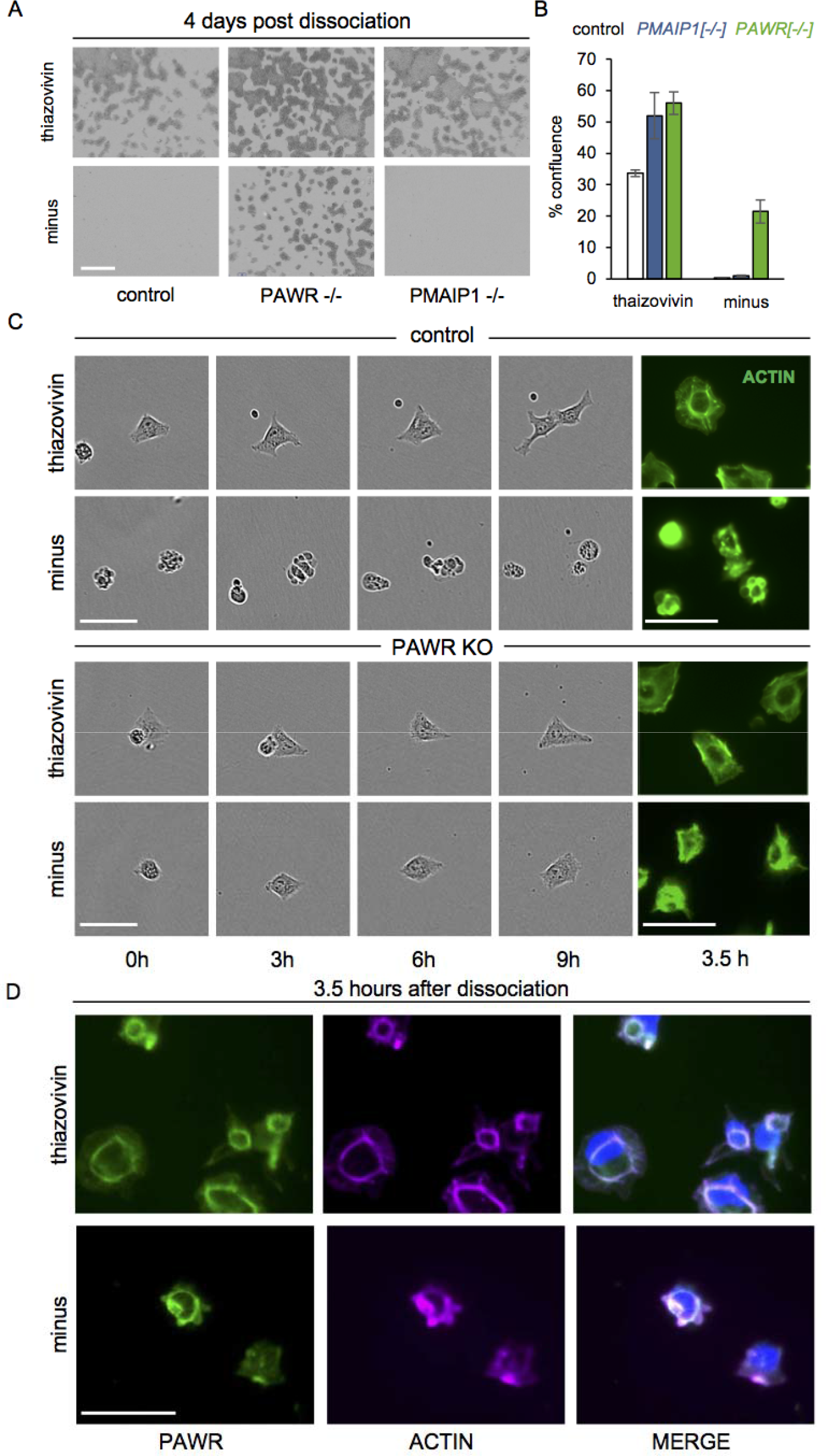
*PAWR* is required for dissociation-induced death. (A) *PAWR* mutants survive single-cell dissociation in the absence of thiazovivin (ROCK inhibitor) treatment. Control cells and *PMAIP1* knockout do not survive without thiazovivin treatment. Bright-field images taken of live iCas9 cells 4 days after dissociation. Scale bar = 800 uM (B) Quantification of survival in the presence or absence of thiazovivin. Percent confluence was measured 4 days after replate in control, *PAWR* knockout and *PMAIP1* knockout cells. Bars represent mean from 3 independent wells with 4 images per well. Error bars +/- 1 std. dev from 4 images per well from 3 independent wells. The dissociation induced survival of *PAWR* mutant hPSCs has been replicated >3 times. (C) Time lapse microscopy of live cells during first 9 hours of replate. Control and *PAWR* knockout hPSCs survive replating in the presence of thiazovivin by extending cellular projections and forming an actin adhesion belt organized with stress fibers. Phallodin stain at 3.5h in fixed cells. Control cells without thiazovivin have abundant membrane blebbing and this is highlighted by the presence of small circular actin rings in phallodin stained cells. *PAWR* mutants have reduced blebbing and an intermediate phallodin staining without small actin rings. Scale bar = 50 uM (D) Immunofluorescence detects PAWR protein colocalizing with F-ACTIN. Following dissociation PAWR protein colocalizes with F-ACTIN in the presence of thiazovivin in adhesion belt like structures and in the absence of thiazovivin in membrane blebs. In green PAWR protein. In magenta phallodin stained F-actin. In blue DAPI stained nuclei. Scale bar = 50 uM

**Figure 6:**
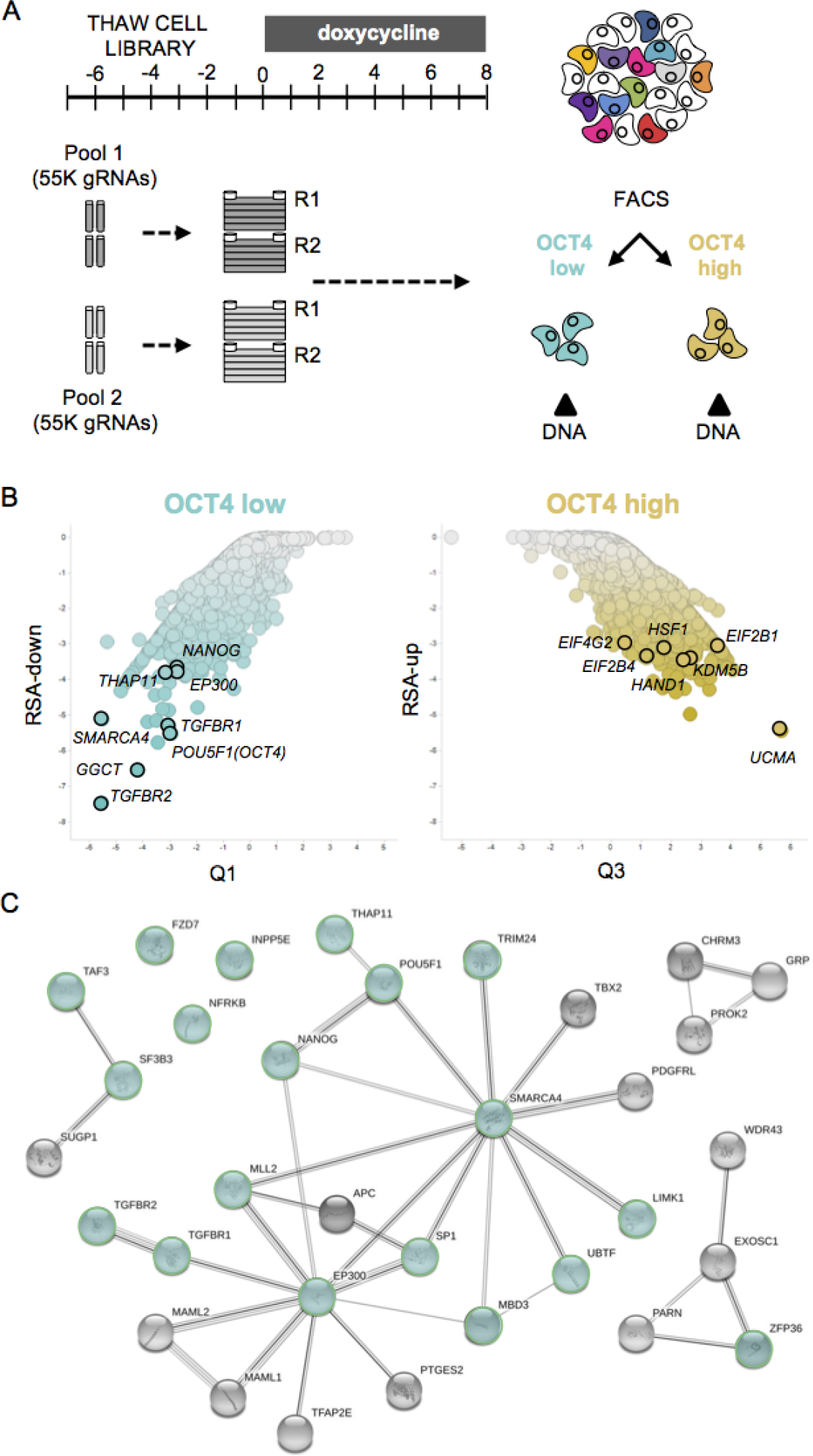
OCT4 FACS-based screen identifies pluripotency gene network. (A) Diagram depicting FACS-based CRISPR screen using an OCT4 specific antibody to sort OCT4 high- and low-expressing cells. Cells were mutagenized with Cas9 for 8 days prior to FACS sorting, DNA isolation and NGS was used to identify an enrichment of sgRNAs in the high and low OCT4 populations. (B) Scatter plot depict gene level results for genome-scale OCT4 FACS screen. Green depict OCT4^LOW^ and gold depicts OCT4^HIGH^ enriched sgRNAs. Y-axis is a p-value generated from RSA down/up analysis. X-axis marks the Z-score (Q1/Q3). (C) String-DB analysis identifies a 20-gene network connected to *OCT4* among gene sgRNAs that were enrich in cells with low OCT4 protein. Genes with published roles in pluripotency, reprogramming and embryo development are highlighted in green.

**Figure 7:**
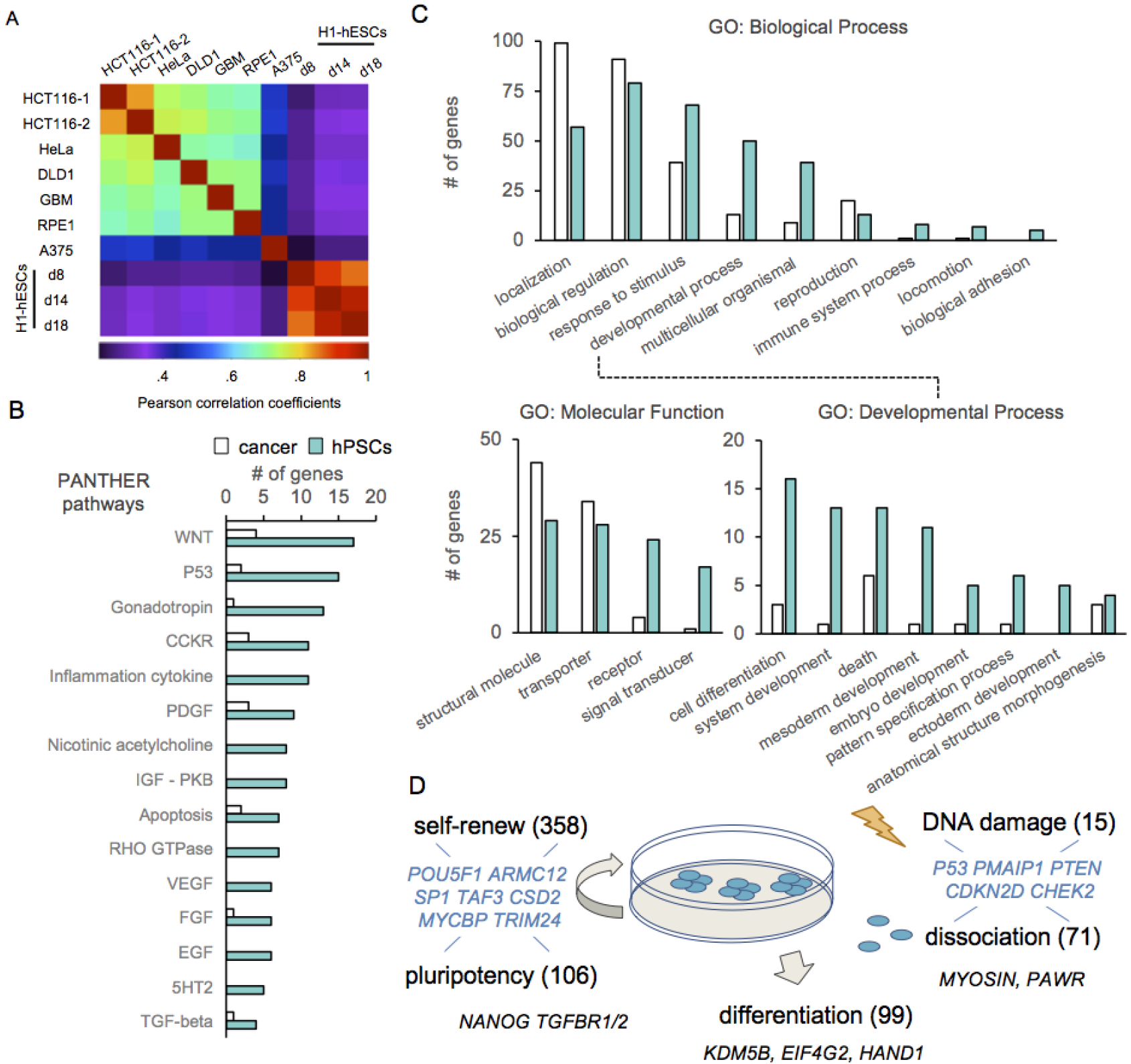
Identification of hPSC-specific essential gene networks. (A) Heatmap of Pearson correlation coefficients adapted from Hart et al., 2015 to include CRISPR screening data in H1-hESCs (B) PANTHER pathway analysis identified 92 enriched pathways in hPSCs. A subset of 15 hPSC-specific pathways are depicted. (C) Depiction of gene ontology categories including biological processes, molecular functions and developmental processes that are specific to hPSCs but not cancer cell lines. (D) Schematic of genes identified by CRISPR screening in hPSCs and their putative functions. 770 fitness genes regulate the self-renewing potential of hPSCs. 113 genes with low OCT4 protein are implicated in pluripotency, 99 genes with increased OCT4 protein may promote differentiation. 20 genes are implicated in the toxic response to DNA damage. 76 genes are implicated in the sensitivity of hPSCs to single-cell dissociation. (B-D) Bioinformatic analysis of 829 core essentials cancer (white bars - Hart et. al., 2015) and 653 hPSC-specific genes (green bars). Genes in multiple categories are shown in blue.

## ACKNOWLEDGEMENTS

We would like to thank R. Maher, J. Alford, S. An, and D. Shkoza for their support with sgRNA cloning. We would like to thank F. Cong, W. Forrester and L. Murphy for access to *PMAIP1* pooled screening results in cancer cell lines. We thank C. Ye for supporting indel analysis and pooled CRISPR screening.

## AUTHOR CONTRIBUTIONS

R.J.I. and A.K designed all experiments and wrote the manuscript. M.R.S, K.A.W and R.D revised manuscript. R.J.I. conducted live cell imaging, immunofluorescence, qPCR and Western blotting S.K. and K.A.W. generated Ngn2-inducible cell lines and differentiated them into Ngn2 neurons. M.S. helped with genome-scale hPSC culture, packaged lentiviral sgRNAs and generated CRISPRi lines. E. F. provided lentiviral supernatants and prepped DNA samples for NGS. J. R. sequenced CRISPR indels. B.A. generated Precision/Recall values using the BAGEL algorithm. S.K. generated RNA expression data in hPSCs and NGN2 neurons and it was analyzed by R.R. G.R.H., Z.Y., and G.M. helped design pooled screen and performed analysis. J. R-H. generated sgRNA libraries. C.R. sequenced pooled fitness screen samples. S.P. sorted OCT4 high and low expressing cells by FACS. D.J.H. sequenced OCT4 FACS screen.

## SUPPORTING INFORMATION

**Figure S1:**
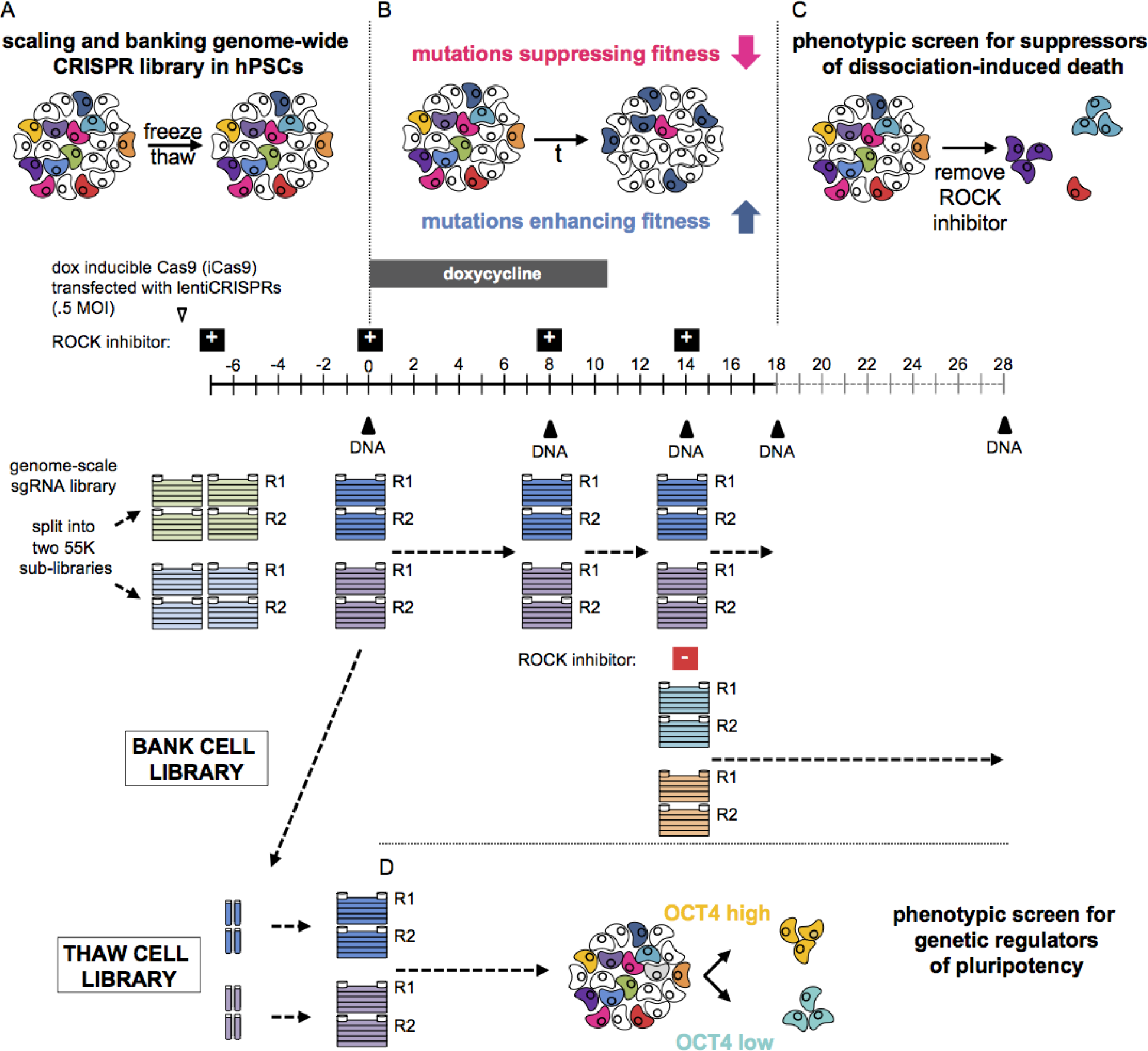
Genome-scale screening in hPSCs. Paradigm for genome-scale pooled screen in hPSCs. (A) hPSCs were transduced with 110,000 lentiCRISPRs in 5-layer CellSTACKs. Following expansion and hPSC CRISPR library banking cells were subjected to subsequent screens depicted in B-D. (B) Fitness screen in hPSCs. Cells were exposed to Cas9 (+dox) for a total of 12 days to ensure complete gene disruption. DNA samples were taken at 0, 8, 14 and 18 days after exposure to Cas9. (C) Positive selection screen for suppressor of dissociation induced cell death. At the end of the fitness screen the genome-scale hPSCs mutant CRISPR library was replated in the absence of ROCK inhibitors (- thiazovivin). After two weeks cells that suppressed dissociation-induced death were collected for DNA isolation. (D) FACS-Based CRISPR screen for regulators of OCT4 protein expression. hPSC CRISPR cell library was thawed and exposed to Cas9 (+ dox) for 8 days before cell sorting to select for cells with high or low expression of OCT4 protein.

**Figure S2:**
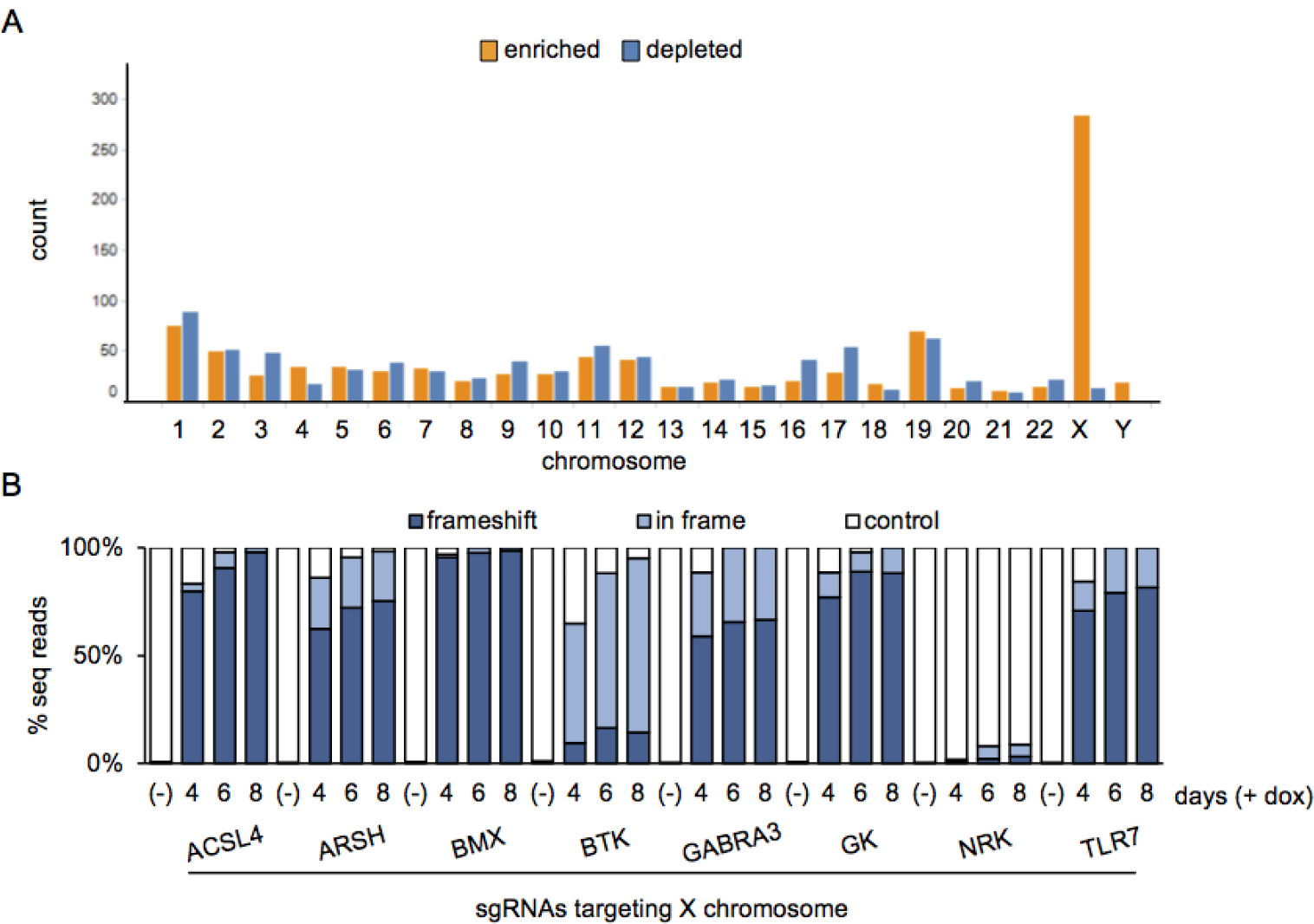
Enrichment of sgRNAs on X and Y chromosomes. (A) Bar chart shows the chromosomal distribution of the top 770 depleted (Files S3) and 946 enriched (RSA-up < −2.25, File S6. (B) NGS quantification indels induced by sgRNAs targeting the X chromosome. Control reads are represented by white bars, in-frame mutations by light blue bars and frameshift mutations by dark blue bars. n=1 DNA sample per sgRNA and cell line. >20,000 sequencing reads per sample.

**Figure S3:**
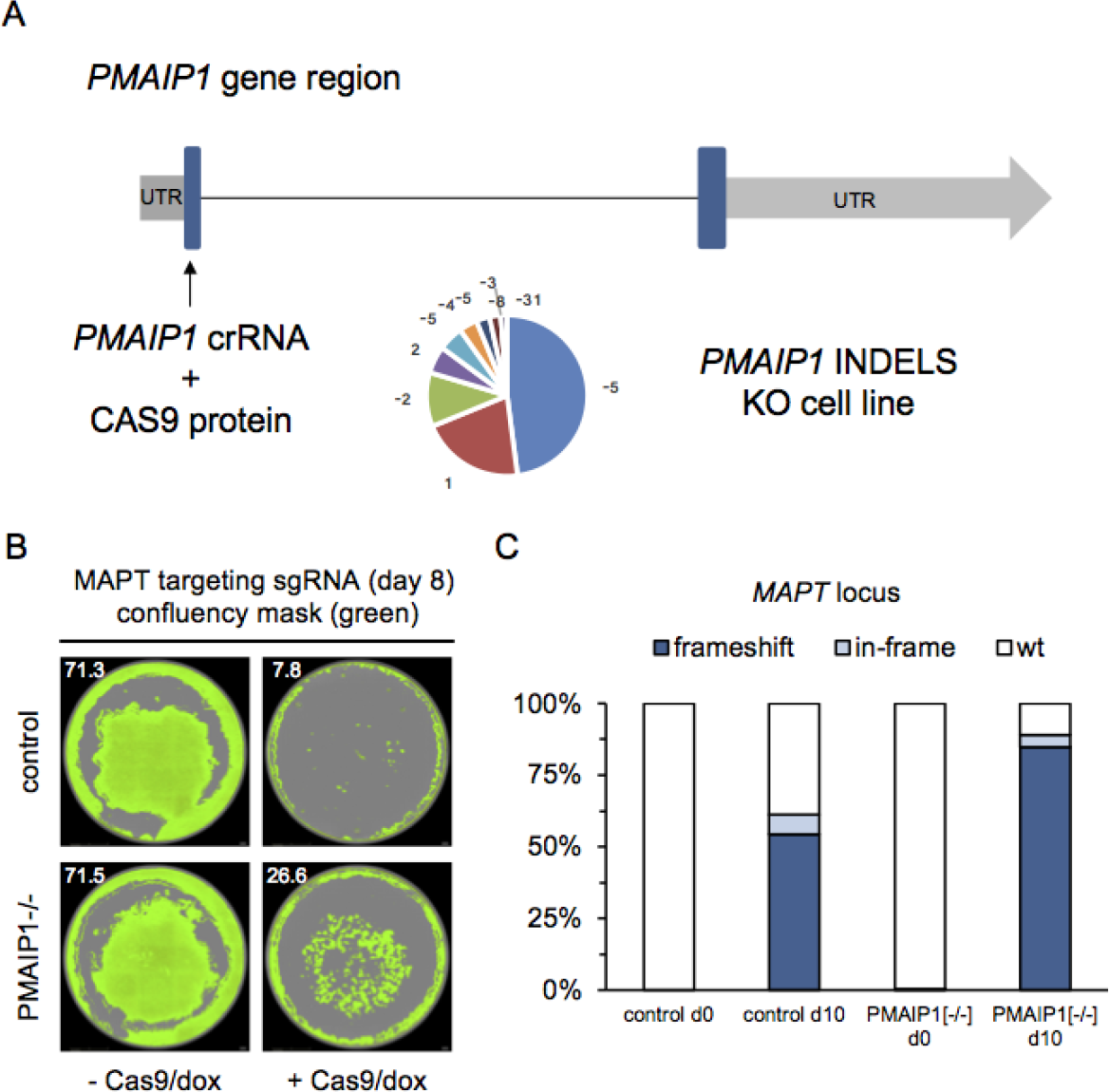
*PMAIP1* knock out hPSCs. (A) iCas9 H1-hESCs were treated with Cas9 RNPs targeting *PMAIP1.* NGS analysis demonstrated that complete knockout of *PMAIP1* was achieved with a spectrum of frameshift indels disrupting *PMAIP1.* n=1 sample, 2627 sequencing reads (B) *PMAIP1* mutations suppress DSB induced toxicity. Whole well images from 24-well plates during editing with *MAPT* sgRNA in H1-iCas9 control and PMAIP1 mutant backgrounds. Confluency mask in green. (C) DNA isolated after 10 of doxycycline treatment shows that on-target *MAPT* editing efficiency is similar between control and *PMAIP1* mutant pools. Control reads are represented by white bars, in-frame mutations by light blue bars and frameshift mutations by dark blue bars. n=1 day 0 samples, n= 2 day 10 samples, >20,000 sequencing reads per sample.

**Figure S4:**
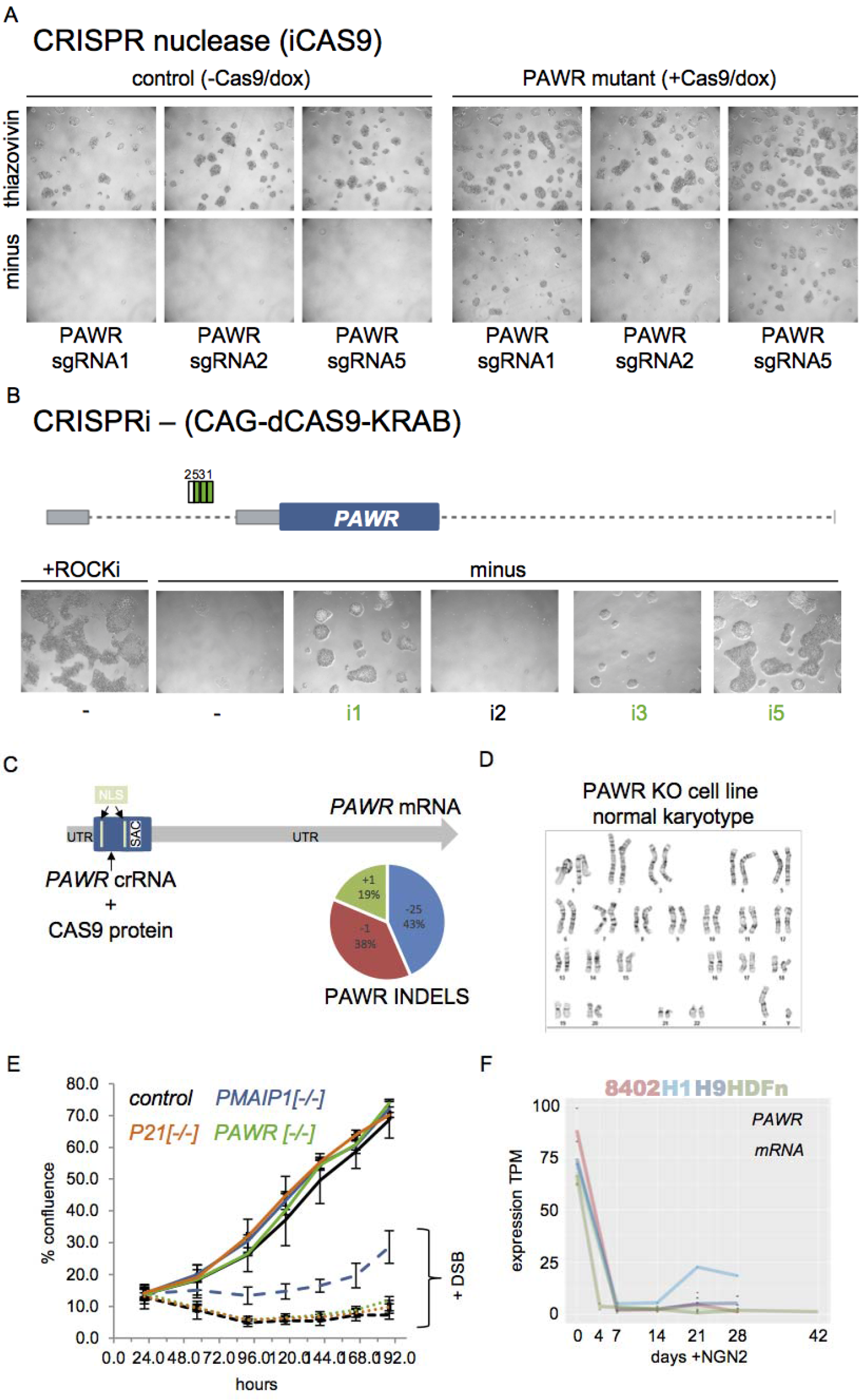
*PAWR* is required for dissociation-induced death. (A) iCas9 expressing H1-hESCs were infected with lentiCRISPRs targeting *PAWR.* PAWR mutant pools were created by exposing cells to Cas9/dox for one week. After mutagenesis, control and mutant cells were dissociated using accutase and replated plus or minus thiazovivin. Control cells die without thiazovivin while PAWR mutants survive. Images were taken 4 days after dissociation. (B) H1-hESC constitutively expressing dCas9-KRAB were infected with CRISPRi sgRNAs targeting the promoter of *PAWR.* 3 out of 4 sgRNAs (i1, i2, i3, i5) tested were able to survive a replate in the absence of thiazovivin while controls died. Images were taken 6 days after dissociation. (C) iCas9 H1-hESCs were treated with Cas9 RNPs targeting *PAWR.* NGS analysis demonstrated that complete knockout of *PAWR* was achieved with a spectrum of frameshift indels disrupting *PAWR.* n=1 samples, 2621 sequencing reads (D) Karyotype analysis revealed that *PAWR* mutants retain a normal Karyotype. (E) *PAWR* mutants do not suppress DNA damage-induced cell death. Live imaging of confluence in *MAPT* sgRNA expressing iCas9 cells +/− DSB (+dox/Cas9) in control, *PMAIP1*knockout cells, and *PAWR*^A^ knockout cells. *PAWR*^−/−^, *P21*^−/−^ knockout and control hPSCs die upon DSB induction while *PMAIP1* mutants survive. Black lines indicate control, blue lines indicate *PMAIP1*mutants, green lines indicate *PAWR*^−/−^ mutants, orange lines indicate *P21*^−/−^ mutants. Solid lines are without dox and dashed lines are cultured with Cas9 (+dox). Y-axis is percent confluency each point represents mean of 3 independent whole-well images. error bars +/− 1 std. X-axis time in days of treatment. (A) *PAWR* is highly expressed in hESCs and iPSCs. Y-axis represents expression in Transcripts Per Kilobase Million (TPM). H1-hESCs (n=1), H9-hESCs (n=3), 8402-iPSCs (n=3) and HDFn-iPSCs (n=4). X-axis represents days after induction of a doxycycline inducible NGN2 expression cassette.

**Figure S5:**
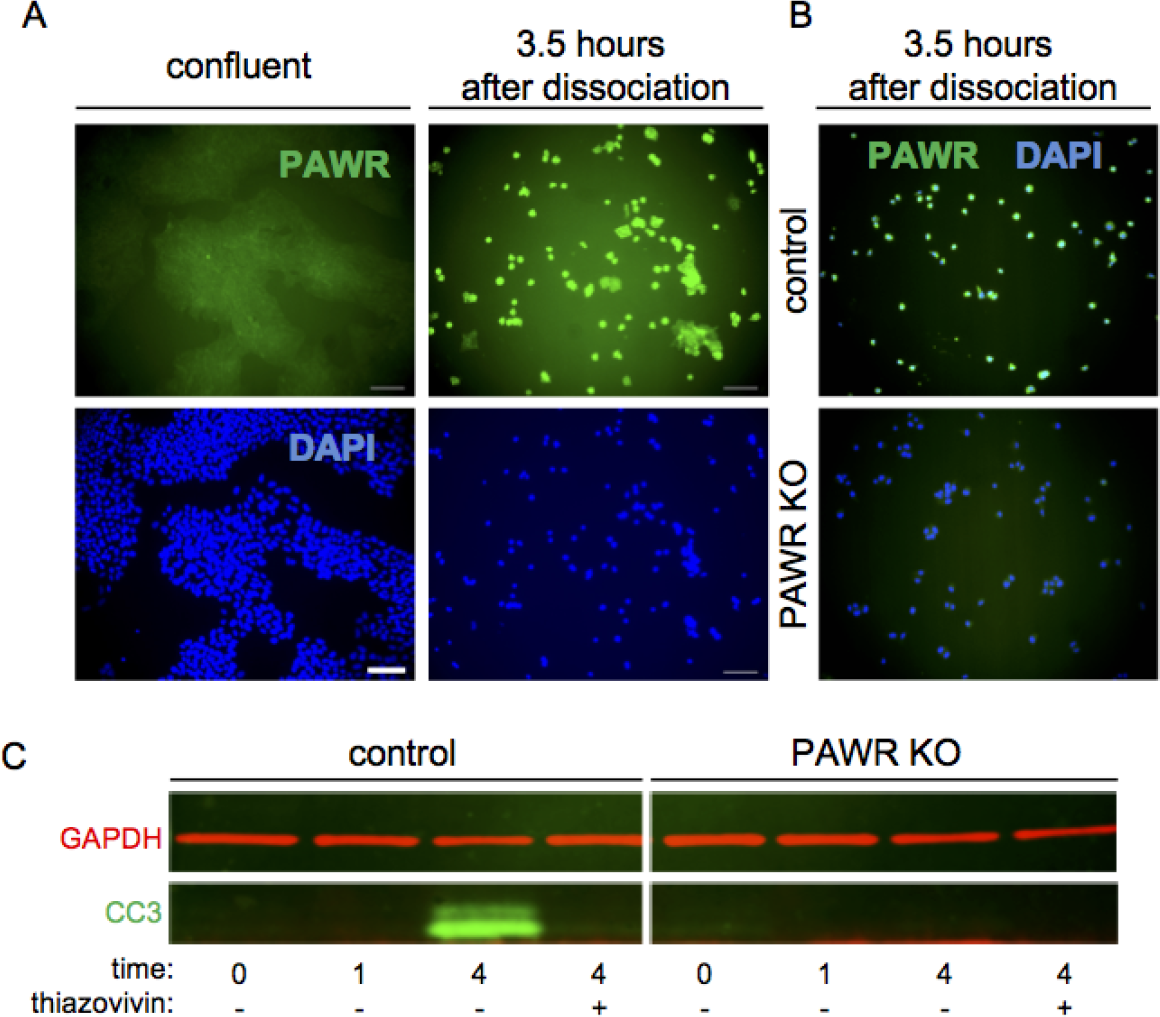
PAWR is induced after dissociation and required for caspase activation. (A) Immunofluorescence using PAWR antibodies detects protein after dissociation but not in confluent colonies. PAWR protein in green. DAPI stained nuclei in blue. Scale bar = 100 uM (B) Immunofluorescence using PAWR antibodies detects protein after dissociation in controls but not *PAWR* knockout cells. (C) In control cells western blot detects caspase activation 4 hours after dissociation in the absence of thiazovivin. *PAWR* mutants do not exhibit caspase activation. In green anti-rabbit Cleaved Caspase-3 (CC3, 17 and 19 kDa) antibodies. In red anti-mouse GAPDH (37 kDa) loading control.

**Figure S6:**
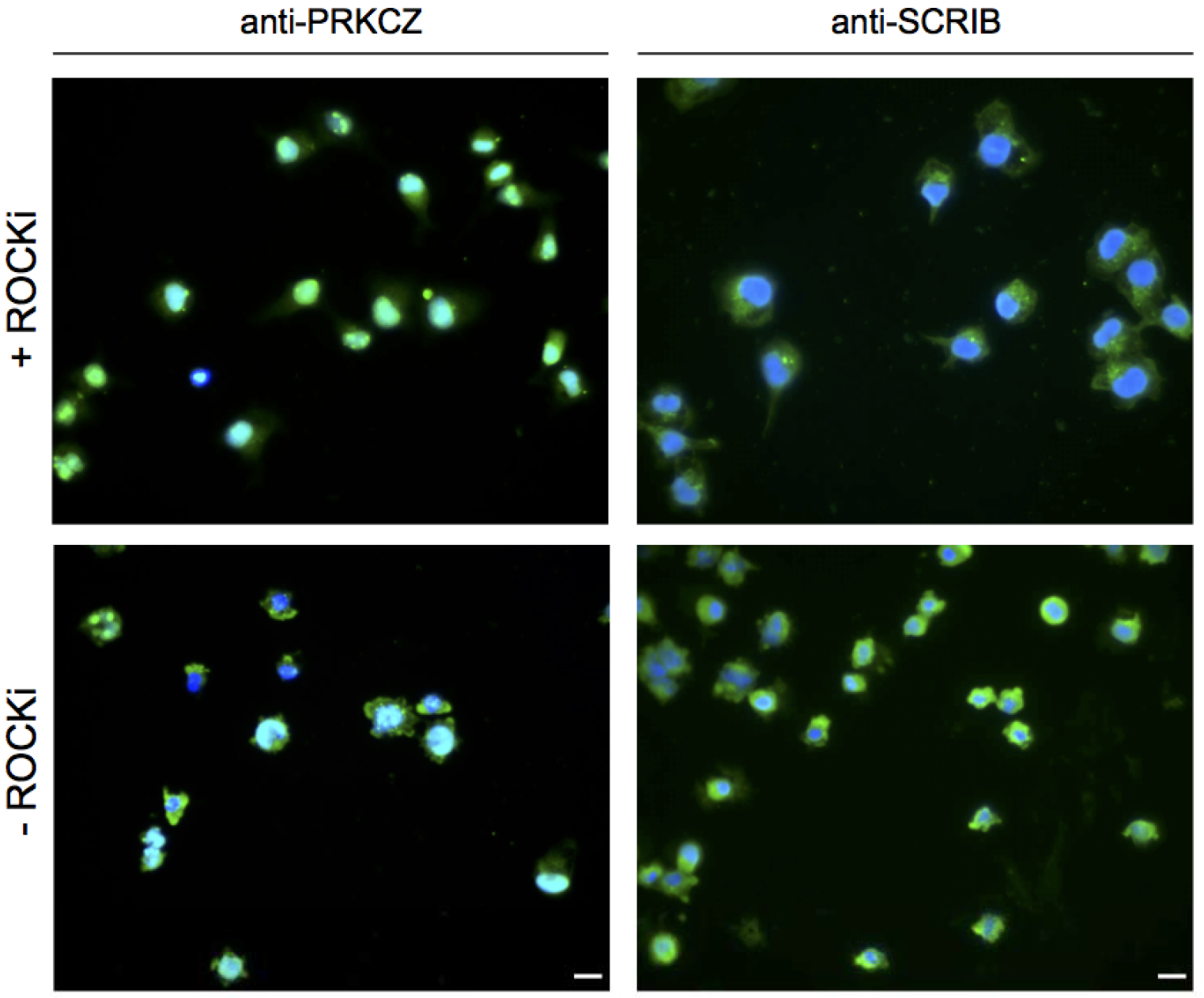
PRKCZ and SCRIB proteins are expressed in dissociated hPSCs. (A) Immunofluorescence detects PRKCZ and SCRIB proteins localizing to membrane blebs in the absence of thiazovivin. In the presence of thiazovivin PRKCZ and SCRIB demonstrate a nuclear or perinuclear localization respectively. In green PRKCZ and SCRIB proteins. In blue DAPI stained nuclei. White scale bar = 20 uM.

## MATERIALS AND METHODS

### Cell lines

H1-hESCs (WA01-NIHhESC-10-0043) and H9-hESCs (NIHhESC-10-0062) were obtained from WiCell. 8402-iPSCs originated from GW08402 fibroblasts from the Coriell Institute and reprogrammed as described by Sun et al., 2016(Sun et al., 2016). hDFN-iPSCs were generated as described by Bidinosti et. al., 2016 (Bidinosti et al., 2016). hPSC lines were free of Myoplasma and tested using the Mycoalert Detection kit (Lonza). SNP fingerprinting confirmed the identify of hPSC lines used. Karyotyping was performed by Cell Line Genetics (Madison,WI).

### Genome Engineering

H1-hESCs expressing dox inducible (i) iCas9 were generated by Ihry et. al., 2018. Dox treated H1-iCas9 cells were subjected to three successive rounds of RNAimax delivery of the two-component synthetic crRNA/tracrRNA (IDT) pairs targeting *PMAIP1, PAWR* and *TP53* as described by Ihry et. al., 2018. *PMAIP1,* and *PAWR* were transfected with a single sgRNA while *TP53* was co-transfected with 3 crRNAs. CRISPR indel analysis detected effcient gene disruption for all three genes and was performed as described by Ihry et. al., 2018.

PMAIP1 crRNA 3 TCGAGTGTGCTACTCAACTC

PAWR crRNA 5 CGAGCT CAACAACAACCT CC

TP53 crRNA 1 GAAGGGACAGAAGATGACAG

TP53 crRNA 2 GAAGGGACAGAAGAT GACAG

TP53 crRNA 4 GAGCGCTGCTCAGATAGCGA

P21 crRNA 1 AAT GGCGGGCT GCATCCAGG

P21 crRNA 4 TCCACTGGGCCGAAGAGGCGG

P21 crRNA 6 GGCGCCATGTCAGAACCGGC

### Pooled Mutagenesis (CRISPR nuclease/interference)

H1-hESCs expressing a constitutive (c) Cas9-KRAB knocked-in to the AAVS1 locus were generated as described by Ihry et., al., 2018. The following lentiCRISPR were used to transduce iCas9 or cCas9-KRAB cells to generate mutant cell pools.

#### CRISPR nuclease

PAWR sgRNA 1 TTTGGGAATATGGCGACCGG

PAWR sgRNA 2 GGTGGCTACCGGACCAGCAG

PAWR sgRNA 5 CGAGCT CAACAACAACCT CC

MAPT sgRNA 1 GAAGT GAT GGAAGAT CACGC

ACSL4 sgRNA TGGTAGTGGACTCACTGCAC

ARSH sgRNA GCAGCACCGT GGCT ACCGCA

BMXsgRNA ATGAAGAGAGCCGAAGTCAG

BTK sgRNA GGAAT CT GT CTTT CT GGAGG

GABRA3sgRNA AAGGACTGACCTCCAAGCCC

GK sgRNA TAGAAAGCTGGGGCCTTGGA

NRK sgRNA CGCCTTCCTATTTCAGGTAA

TLR7 sgRNA CAGTCTGTGAAAGGACGCTG

#### CRISPR interference

PAWR sgRNA 1 GGCGCGCTCGAGGACTCCAA

PAWR sgRNA 2 GTTGCAGGGTGGGGACCCGG

PAWR sgRNA 3 GCT GGCCGGT AGT GACT GGT

PAWR sgRNA 5 GGCT GCT GGCCGGT AGT GAC

### Large-scale culture of hPSC

H1-hESCs with AAVS1 knock in of the iCas9 transgene were generated and cultured in TeSR-E8 media (STEMCELL TECH.-05940) on vitronectin (Gibco-A 14700) coated plates as described by Ihry et. al., 2018. Pilot studies with a sub-genome sgRNA library used a total of 40 individual T225 flasks and was cumbersome (Ihry et. al., 2017). Daily feeding and passaging in which multiple flask needed to be pooled was time consuming and increased the risk for contamination. To minimize the manipulation required during feeding and passaging we used 5-layer CellSTACKs. The vessels are large enough to contain an entire 55,000 sgRNA at roughly 1000x coverage per sgRNA at a seeding density of 21,000 cells/cm[2] (Seed 66 million cells for ~1200x). In practical terms only 4 to 8 5-layer CellSTACKs were growing at a given time during the month-long genome-scale CRISPR screen (Fig. S1). Given the expense of E8 media Penicillin-Streptomycin (pen/strep) at 100 U/mL was added as an additional insurance policy for the first screen at scale. After running a lengthy genome-scale screen and becoming experienced with large-scale hPSC culture it is possible to run future screens without pen/strep (Fig. S2).

### lentiCRISPR packaging

For a one-layer CellSTACK 42 million HEK293T (66,000 cells/cm^2^) were plated in 100 mL of media (DMEM +10% FBS + 1x NEAA, no pen/strep). One day after seeding, cells were transfected with single lentiCRISPR plasmids in 6-well plates or pooled lentiCRISPR plasmids in CellSTACKs. For a one-layer CellSTACK 102 uL of room temp TransIT (Mirus MIR 2700) and 3680 uL Opti-MEM (Invitrogen 11058021) were mixed incubated in a glass bottle for 5 minutes at room temp. 94.5 ug of Lentiviral Packaging Plasmid Mix (Cellecta CPCP-K2A) and 75.6 ug of the lentiCRISPR plasmid library was added to the transfection mix and incubated for 15 minutes at room temp. After incubation, the mix was added to 100 mL of fresh media and the cells were fed. The next day the transfected cells received 100 ml fresh media. After 3 days of viral production supernatants were filtered (.45uM corning 430516) and aliquoted in to 1ml tubes for storage at −80 C.

### Large-scale transduction of hPSC

We conducted a genome-scale CRISPR screen using a 110K sgRNA library (~5 sgRNAs per gene, split into two 55K sgRNA sub-pools, DeJesus et al., 2016). We desired 1000x coverage of each sgRNA to offset cell loss from double strand break (DSB)-induced toxicity (Ihry et. al., 2017). To screen a sufficient number of cells we infected hPSCs with lentiCRISPRs in 5-layer CellSTACKs. Cells were infected at .5 MOI to ensure each cell was infected with no more than a single sgRNA. After puromycin selection cells were expanded for one week without dox to be pelleted for DNA, banked or screened (Fig. 1A, S1).

Pooled CRISPR screens rely on cells being efficiently transduced at less than or equal to .5 MOI. We developed a reverse transfection method for hPSCs without polybrene that resulted in an efficient transduction with low volume exposure to HEK293T lentiviral supernatants. LentiCRISPR plasmids expressed a constitutive RFP and puromycin resistance to mark and select for infected cells respectively. Viral titer of the two 55,000 sgRNA libraries expressing RFP in 6-well plates determined that less than 25 uL in 1.5mL of media was required for .5 MOI. These calculations scaled appropriately in 5-layer cell stacks with 500 mL of media and approximately 50% of the cells were RFP positive in the absence of puromycin.

A genome-scale lentiCRISPR library targeting each gene 5 times has been split into 55,000 sub-pools (pool 1 and 2). To screen at 1000x per sgRNA, 264 million hPSCs are infected in 4x 5-layer CellSTACKS (12,720 cm^2^, 21,000 cells/ cm^2^). Cells are infected at .5 MOI to ensure only one sgRNA is expressed per cell (+sgRNA/puro^R^/RFP). After puromycin selection cells are expanded until confluent (4-6 days). At this point 40 million cells (10 million/ml) can be banked in 5ml cryovials for use at a later date.

### Banking lentiCRISPR infected hPSC library

One 5-layer cell stack was treated with 200 mL accutase that was evenly distributed among layers. After incubation at 37 C for 10 minutes, accutase (Gibco-A1110501) was neutralized with 200 mL E8 media. Cells were counted and pelleted to be resuspended at a concentration of 10 million cells per ml in a solution of 40% Tet-free FBS (Seradigm #1500-500) and 10% DMSO (Sigma D2650) and 50% E8 media. 4 ml aliquots were placed in 5 ml cryovials and frozen in a Mr. Frosty (Thermo Scientific 5100-0050) at −80 C overnight before long-term storage in liquid nitrogen. Thawing of cells banked in 5ml cryovials showed an average viability around 85% for both lentiCRISPR pools. Viability was assayed using a Nexcelom Cellometer Auto 2000 and AO/PI (Nexcelom CS2-0106) staining solution.

We tested the effect of freezing and thawing on the representation of the sgRNAs in the library by thawing the cell library at either 700x (40 million cells per 55k sub-pool) or 2100x (120 million cells per 55k sub-pool) cells per sgRNA. Cells were thawed in a 37 C water bath and transferred to a 50mL conical with E8 media and centrifuged at 300 g for 3 minutes. Pelleted cells were resuspended in E8 media and replated at a density of 30 to 40*10^6^ cells/cm^2^. 3 cryovials with 120 million cells was thawed and plated on a 5-layer cell stack (120,000,000 /55,000 = 2100x).

After thawing and feeding the cells for two days, DNA was isolated and analyzed by NGS to measure the representation of each sgRNA in the pool. Each sample was compared to the original lentiCRISPR plasmid library to generate a log_2_(fold change) value. Both day zero and freeze/thaw samples had an over 87% alignment of sequencing reads and fewer than 25 missing barcodes per replicate. Calculating the log2(fold change) revealed no significant change in the representation of sgRNAs before (day 0) or after one freeze/thaw cycle. Pearson correlation analysis demonstrated there was a high correlation between the day zero samples and the freeze/thaw samples (Fig. 1B).

### OCT4 FACS-BASED screen

Cells were dissociated using accutase for 10 min at 37 C to create a single-cell suspension which was strained using a 40-micron filter and was counted. After removing accutase, pelleted cells were resuspended in a volume of 1 million cells/mL for the staining protocol. For each replicate 55 million unsorted cells were frozen down prior to fixation. The remaining cells were fixed in 4% PFA in PBS for 10 minutes at room temperature on a rocker. Cells were spun down at 300 RCF for 3 min between each subsequent solution change. Cells were washed with .1% Triton-X in PBS after fixation and blocked in 2% goat serum, .01% BSA and .1% triton X in PBS for 1hr at room temperature. Conjugated (AF488) primary antibodies specific to OCT4 (CST 5177) were diluted in blocking solution (1:200) and incubated with cells on a rocker over night at 4 C. Prior to FACS analysis cells were resuspended in PBS at a concentration of 30 million cells/mL. A total of 1.2 billion cells were sorted into OCT4^low^ (50 million cells) and OCT4^high^ (61 million cells) populations using an ARIA III (BD).

### DNA isolation

For each replicate, 55 million cells (1000x) were pelleted and genomic DNA was isolated using QIAamp DNA Blood Maxi Kit (Qiagen 51194) as directed by manufacturer. Isolating genomic DNA from 4% PFA fixed cells was performed by utilizing phenol chloroform extraction. Cells were resuspended in 500ul TNES (10mM Tris-Cl ph 8.0, 100mM NaCL, 1mM EDTA, 1% SDS) and incubated overnight at 65 C. After allowing the samples to cool, 10ul of RNase A (Qiagen 19101) and samples were incubated at 37 C for 30 minutes. Next, 10ul of proteinase K (Qiagen 19133) and incubated for 1 hour at 45 C. Add 500ul of PCIA (Phenol;Cholorform;Isoamyl alcohol ph 8) (Thermo 17908) and vortex. Spin samples in a centrifuge at max speed for 2 minutes. Transfer the aqueous phase to 500ul of PCIA and vortex. Spin at max speed for 2 minutes. Transfer the aqueous phase to 450ul of chloroform and vortex. Spin at max speed for 2 minutes. Transfer aqueous phase to 40ul of 3M NaAcO ph 5.2. Add 1 ml of 100% ethanol, mix and precipitate DNA for 1 hour on ice. Spin at max speed for 2 minutes. Decant and wash with 1ml 70% ethanol. Spin at max speed for 1 minutes. Decant and air dry the pellet. Resuspend in 50ul of nuclease free H_2_0. mRNA expression qPCR and RNA-seq analysis were performed as described by Ihry et. al., 2018. Median TPM values for H1-hESCs are available upon request. RNA-seq data for in 4 iPSC control lines subjected to NGN2 differentiation is available upon request but is restricted to the expression for *PAWR* and *PMAIP1.*

### Assaying sensitivity to DNA damage

Control and mutant iCas9 H1-hESCs expressing a *MAPT* targeting sgRNA were monitored daily post-media change using an IncuCyte zoom (Essen Biosciences). At the onset of the experiment cells were plated at density of 10,500 to 21,000 cells/cm^2^ and cultured plus or minus dox for the duration. Confluence was calculated using the processing analysis tool (IncuCyte Zoom Software).

### Assaying survival without ROCK inhibitor

Control and mutant iCas9 H1-hESCs were dissociated with accutase for 10 minutes. Flowmi 40- micron cell strainers (BEL-ART H13680-0040) were used to ensure a uniform single-cell suspension prior to replating cells. Cells were plated at a density of 10,500 to 21,000 cells/cm^2^ plus or minus thiazovivin (ROCK inhibitor). Timelapse images were taken in 3 hr intervals using IncuCyte zoom (Essen Biosciences). The confluence processing analysis tool (IncuCyte Zoom Software) calculated confluency for each sample.

### Immunofluorescence and Western Blotting

Immunofluorescence staining of fixed cells was performed as described by Ihry et. al., 2018. Protein lysates were made by vortexing cell pellets in RIPA buffer (Thermo Scientific 89901) supplemented with Halt protease inhibitor cocktail (Thermo Scientific 78430) and phosphatase inhibitors (Thermo Scientific 1862495). Samples were incubated at 4C for 10 minutes before centrifugation at 14,000 x g for 10 minutes at 4C. Supernatants were transferred to new tubes and quantified using the BCA protein assay kit (Thermo Scientific 23225) and a SpectraMax Paradigm (Molecular Devices) plate reader. Samples were prepared with NuPAGE LDS sample buffer 4x (Invitrogen NP0008) and NuPAGE sample reducing agent (Invitrogen NP0009) and heated for 10 minutes at 70 C. Chameleon Duo Pre-stained Protein Ladder (LiCOR P/N 928-60000) was loaded alongside 10 ug of protein per sample on a NuPAGE 4-12% Bis-Tris Protein Gels, 1.5 mm, 10-well (Invitrogen NP0335BOX). Gel electrophoresis was performed at 150 V for 1hr in NuPAGE MOPS SDS Running buffer (20X) (Invitrogen NP0001) using a XCell SureLock Mini-Cell (Thermo Scientific EI0002). Transfer was performed using an iBlot 2 dry blotting system (Thermo IB21002 and IB23002) as described by manufacturer. Blots were blocked in TBS blocking buffer (LiCOR 927-50000) for 1 hour at room temperature. The blots were then incubated with primary antibodies diluted in TBS over night at 4 C. Blots were washed 3X in PBST and incubated with secondary antibodies diluted in TBS for 2 hours at room temperature. Blots were imaged using an Odyssey CLx (LiCOR).

#### Primary antibodies

Phallodin-647 (ThermoFisher Scientific A22287) - 1:40 SCRIB (Abcam ab36708) - 1:100 (IF)

PRKCZ (Abcam ab59364) - 1:100 (IF)

PAWR/PAR-4 (CST-2328) - 1:100 (IF)

Cleaved Caspase-3 Asp175 (CST-9661) - 1:200 (IF) 1:1000 (WB)

GAPDH (Enzo ADI-CSA-335-E) - 1:1000 (WB) aTUB (Sigma 76199) - 1:10000 (WB)

#### Secondary Antibodies

IRDye 800CW anti-rabbit (LiCOR 926-32211) - 1:5000

IRDye 680RD anti-mouse (LiCOR 926-68070) - 1:5000

AF488 conjugate Goat anti-Rabbit IgG (H+L) (ThermoFisher-A-11008) - 1:500

## DATA AVAILABILITY

Raw data and results generated for human pluripotent stem cells is available upon request. File S1 includes log2 fold changes values for each sgRNA across conditions. File S2 and S6 includes RSA-up and -down values for the fitness screen and File S8 has RSA values for the OCT4 FACS screen.

## CODE AVAILABILITY

RNA-seq data processing and pooled CRISPR NGS was conducted using open source software.

